# Low-cost monophasic transcranial magnetic stimulator

**DOI:** 10.64898/2026.08.25.747050

**Authors:** Maxence Lapatrie, Yuzuha Isetani, Jathav Puvirajan, Antonella Catanzaro, Siqi Lyu, Han C. Nguyen, William Mathieu, Milica Popović

**Author notes:** **Corresponding author’s email address**.

## Abstract

Transcranial magnetic stimulation (TMS) excites neurons noninvasively by electromagnetic induction and is used in neurophysiology research and in approved therapy for depression. Commercial stimulators cost tens of thousands of dollars. Existing open-source designs are either low-energy and unvalidated or rely on expensive switches and laboratory infrastructure. We present a monophasic, fixed-pulse-shape TMS device built at a parts cost of ∼USD 700 which, under specific modeling assumptions, can exceed average human motor thresholds. Our design assumes access to basic, off-the-shelf equipment such as a 24 V power supply unit, an oscilloscope, and a few basic tools. The device charges a 230 µF film-capacitor bank and discharges it through a self-wound figure-of-eight coil using a thyristor, producing a fixed pulse with a positive lobe lasting approximately 90 µs. A Zero-Voltage Switching (ZVS) driver-based charging circuit charges the capacitor bank up to 1460 V from a 24 V bench supply. Three galvanically isolated voltage domains, redundant interlocks, and passive and active discharge paths help mitigate the safety risks involved with handling lethal energy levels. We also present a low-cost way to characterize the device by reconstructing coil 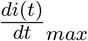 from pickup-coil 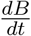 maps to estimate the induced cortical *E*-fields. At the maximum capacitor voltage, the recovered 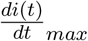 is 110.86 A/µs, giving estimated 99.9th percentile cortical *E*-fields of 159 V/m at Oz and 196 V/m at C3 on an example anatomy. Although not yet approved for clinical trials and routine stimulation, the device demonstrated the possibility of a cost-effective TMS unit.

Graphical abstract
Low-cost open-source monophasic transcranial magnetic stimulator

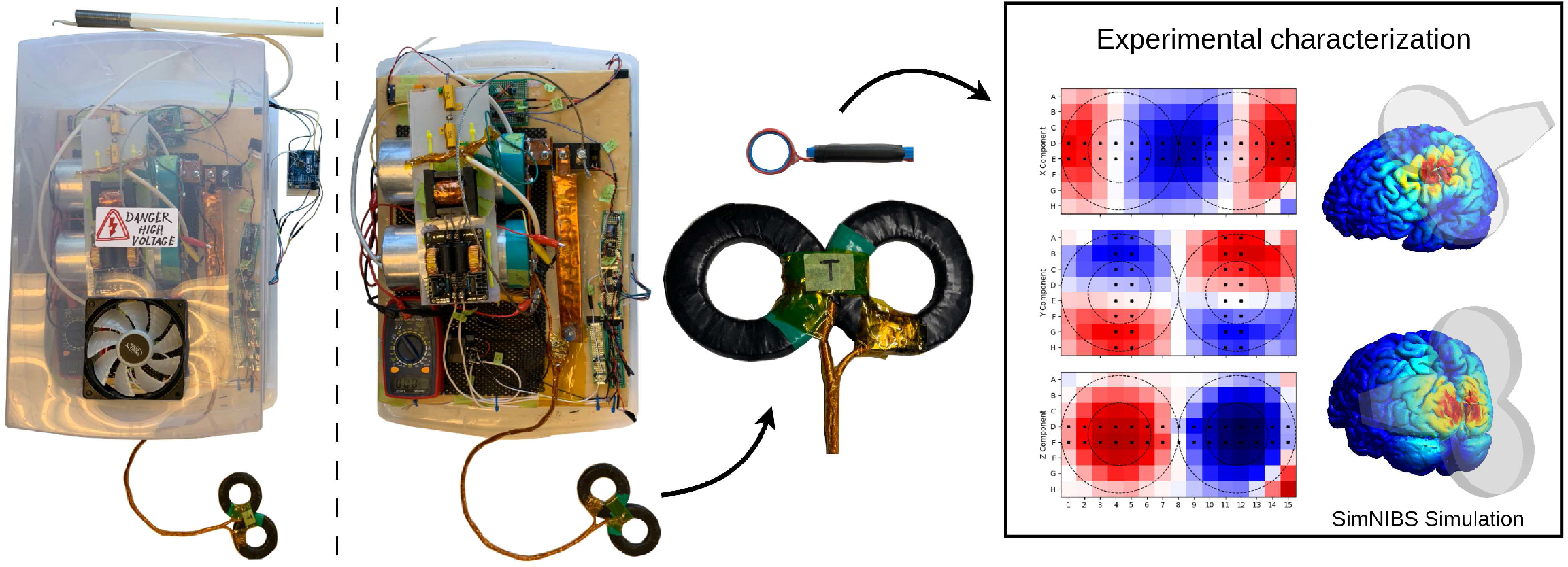

**Highlights:**

- A monophasic TMS was built for USD 657.34 from accessible components.
- The stimulator reaches a peak coil *dI/dt* of 110.86 A/µs.
- Pickup coil field mapping agreed with measurements from a clinical TMS system to within 5.55%.
- Simulated cortical fields reached 159 V/m at Oz and 196 V/m at C3 on an example anatomy.
- Galvanic isolation, redundant interlocks and discharge paths improve safety.

## 1 Hardware in context

Transcranial Magnetic Stimulation (TMS) is a non-invasive brain stimulation method introduced at the University of Sheffield in 1985 [1]. Unlike transcranial electrical stimulation, another non-invasive brain stimulation method which drives currents through scalp electrodes and is limited by the discomfort of high surface current densities, TMS does not require contact with the scalp for effective stimulation. A brief, rapidly changing magnetic field passes through the scalp and skull and induces an electric field directly in the underlying tissue [2].

A capacitor bank is discharged through a coil, placed strategically near the head surface, producing a large 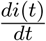. The resulting time-varying magnetic field induces an electric field in the tissue, by Faraday’s law. The component of the induced field directed along an axon creates spatial variations in extracellular potential that polarize the cell membrane and can initiate an action potential. Neural recruitment depends not only on the field magnitude, but also on the pulse waveform, neuronal orientation and morphology, membrane dynamics, and the spatial variation of the field along the axon. Tissue-conductivity boundaries shape the induced electric field, while axonal bends and terminations change its component along the neuron and may create sites of strong membrane polarization [2]. Biological tissues are non-magnetic, their relative magnetic permeability is conventionally approximated as that of free-space (unity) [3]. The generated magnetic fields therefore penetrate the scalp and skull with little attenuation. The induced electric-field distribution is instead shaped primarily by the coil geometry and the distribution of tissue conductivity in the head.

Delivered repetitively, TMS has been shown to induce lasting changes in cortical excitability through forms of synaptic plasticity such as long-term potentiation (LTP) and long-term depression (LTD). TMS devices and protocols are FDA-cleared for major depressive disorder, obsessive-compulsive disorder, anxious depression, smoking cessation, and migraine with aura, with each indication tied to a specific device and stimulation protocol [4]. Despite this established clinical and research role, commercial stimulators remain expensive. A monophasic stimulator comparable to ours, the Magstim 200^2^, costs on the order of tens of thousands of dollars, placing it out of reach for many teaching labs, smaller research groups, and independent builders [5].

This cost barrier has motivated several open and do-it-yourself TMS designs. Among the systems we reviewed, however, none combined all the objectives of the present work, namely: high power output relevant to TMS research, construction from easily obtainable components, operation without specialized power infrastructure, quantitative characterization, and documented procedures for safe assembly, operation, and de-energization. Hobbyist designs generally provide limited validation and little formal safety documentation [6, 7]. Higher-power systems have also been reported, but some require expensive specialized laboratory equipment or costly components, while others lack a complete and current bill of materials or do not report sufficient details to permit a present-day comparison [8, 9, 10, 11].

In this work, we present a monophasic TMS research prototype with low production cost of ∼USD 700, using off-the-shelf components. Its construction and characterization require only a 24 V power supply, a multimeter, an oscilloscope, and common hand tools beyond the components included in the bill of materials. We quantitatively characterize its magnetic fields and, through simulation, show induced cortical *E*-fields in one example head model above published average human motor threshold estimates. The prototype is intended for construction, evaluation, and methodological development by researchers with appropriate high-voltage and pulsed-power expertise. We emphasize that this is not yet an approved medical device, as it has not been validated through clinical trials.

The provided safety layers are intended to reduce specific foreseeable risks and failure modes. They do not make the stored energy safe. Separation of the control, low-voltage power, and high-voltage domains through galvanic and electric isolation reduces the possibility of high-voltage faults or inductive transients propagating into the control electronics and host computer. Series lid, dead-man, and charge-cutoff interlocks reduce the risk of charging under unintended operating conditions, while independent hardware and software overvoltage protection limits overcharging. Bleeder resistors address the possibility of the capacitor bank being inadvertently left charged, and a manual discharge stick allows for almost immediate de-energization if the enclosure needs to be opened. Nevertheless, in our tests, we reached a maximal stored energy in the capacitor bank of around 245 J, which can be lethal if direct contact is made with the bank. Although mindfully assembled, these measures are not a substitute for technical competence, controlled laboratory procedures, and safety certifications.

## 2 Hardware description

- Study high-output monophasic TMS without requiring a commercial stimulator or specialized pulsed-power infrastructure.
- Prototype self-wound stimulation coils and quantify how changes in coil geometry, inductance, and placement affect the generated magnetic field distribution.
- Characterize TMS coils using a low-cost pickup-coil setup for *dB/dt* mapping and use SimNIBS [12] for *E*-field mapping.

This section describes a monophasic, fixed-pulse-shape TMS device developed for circuit, coil, and field-characterization studies in a controlled laboratory setting. The device, shown in Figure 1, stores energy in a 230 µF film-capacitor bank and discharges it through a self-wound figure-of-eight coil using an SCR, producing an effective positive pulse duration of approximately 90 µs. The charging transformer provides nominal voltage taps up to 1875 V. We calculate a theoretical lossless maximal voltage for our capacitor-coil combination of 1428.4 V, and we find a practical limit of 1460 V. When we refer to maximal voltage, we mean the practical 1460 V limit. This corresponds to approximately 245 J of stored energy at maximal tested voltage. At this voltage, field mapping using a pickup coil revealed a coil *dI/dt* of 110.86 A/µs, from which the 99.9th-percentile cortical electric fields were estimated as 159 V/m at Oz and 196 V/m at C3. Testing was limited to intermittent single-pulse operation. At the maximum capacitor-bank voltage of 1460 V, we delivered 30 pulses during each 15 min test period, with an interpulse interval of 30 s. Consecutive test periods were separated by a 15 min cooling interval. Sustained repetitive operation and its thermal limits were not established.

**Figure 1:**
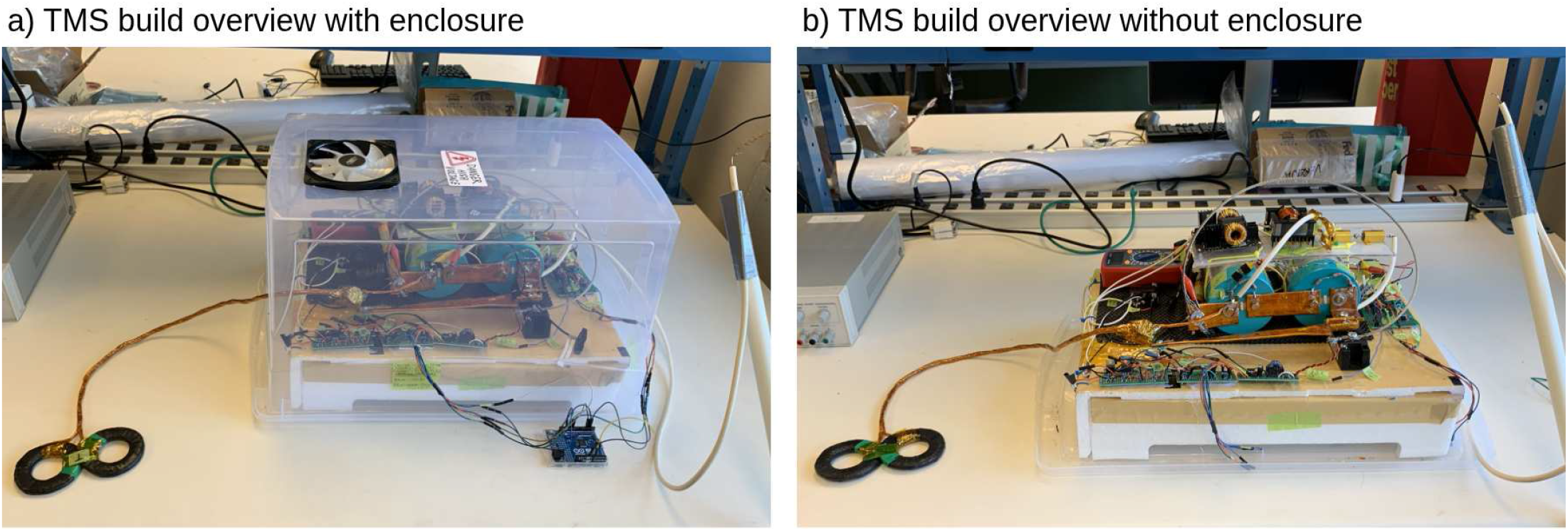
Overview of the TMS. **a) TMS build overview with enclosure**. The TMS components are inside the storage container. On the right side, the safety stick can be seen connected to the CAP-terminal with its other lead left floating. Jumper cables exit the container from holes drilled on the sides and connect to the microcontroller. On the left, the TMS coil exits the front of the enclosure through a hole. **b) TMS build overview without enclosure**. The lower portion of the storage container is removed, exposing the internal components of the TMS device.

The circuit charges a film-capacitor bank to a set voltage, then discharges it through a figure-of-eight coil using a thyristor. The discharge circuit is piecewise. Initially, the circuit behaves as an underdamped system resulting in a positive current pulse in the coil. Then, as the capacitor bank voltage crosses zero, the system becomes overdamped, causing the pulse energy to dissipate in a damping resistor. The pulse shape is fixed by the passive components and is not adjustable during operation. The coil current rises and decays in one direction while the *dI/dt* and induced electric field have a short dominant phase followed by a weaker opposite phase.

The system spans three electrical domains. The control domain holds the host computer and microcontroller. The control domain uses GNDA as common ground, which corresponds to the microcontroller ground. The power domain holds the 24 V supply and all control circuitry. Its ground is labeled GND and is the PSU ground. The high voltage domain holds the capacitor bank and everything electrically continuous with it. This part is highly dangerous and therefore the most isolated part of the circuit. Its ground is labeled CAP- and corresponds to the negative terminal of the capacitor bank. The three domains are galvanically isolated. Logic signals cross into the power and high voltage domains only through optocouplers. A pulse transformer isolates the gate driver control subcircuit from the high-voltage SCR. Similarly, the charging circuit uses a step-up transformer to charge the capacitor bank.

The device produces an effective pulse length of around 90 µs. For this monophasic device, we use the term effective pulse length to refer to the rise time to peak current. This is the duration of the first quarter-lobe, during which the current rises, or analogously, during which the capacitor voltage falls from its peak. The pulse length is set by the inductance and capacitance of the circuit. In commercial monophasic devices, the effective pulse length is often around 70-100 µs [13, 14]. Since the current waveform is difficult to obtain, we instead measure the capacitor voltage during discharge and derive the key system properties from it. Figure 2 displays those key definitions on an example waveform.

**Figure 2:**
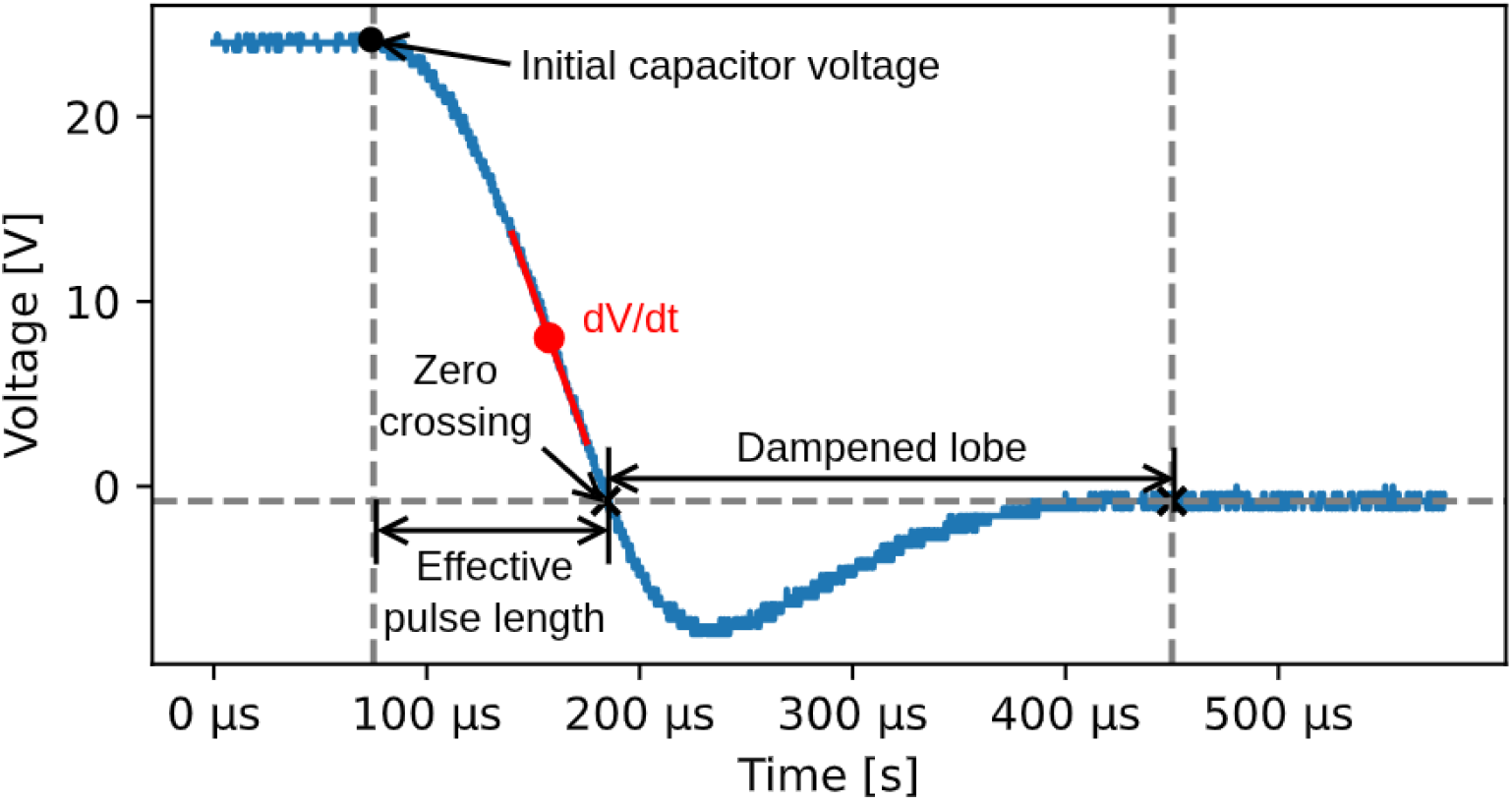
Capacitor discharge waveform and key definitions. The blue trace corresponds to the voltage across the capacitor bank during a pulse discharge with an initial bank voltage of 24 V. The 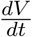 corresponds to the slope of the voltage function. The time between the beginning of the discharge and the first zero crossing is called the effective pulse length. After the first zero crossing, the system becomes overdamped. We call this lobe the dampened lobe.

After the first quarter lobe, the system becomes overdamped and generally takes around a millisecond to reach its equilibrium. The strong 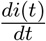 in the coil during the first quarter followed by a weak 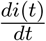 for the remainder of the pulse is what defines monophasic pulses and makes them directionally selective.

The effective pulse length must be selected relative to the chronaxie of the targeted cortical neurons. Chronaxie is a strength–duration parameter that characterizes how briefly a stimulus can act while still reaching the activation threshold. The threshold stimulus strength rises as the pulse is shortened. In the exponential (Lapicque) form derived from a first-order RC membrane model [15],

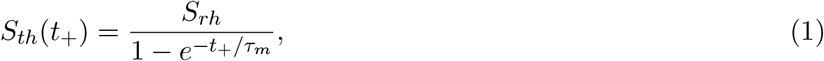

where *S*_*th*_ is the threshold strength, *S*_*rh*_ the rheobase (the threshold for an infinitely long pulse), *τ*_*m*_ the neuronal membrane time constant, and *t*_*c*_ the chronaxie. The chronaxie is the pulse duration at which the threshold equals twice the rheobase. Substituting *S*_*th*_ = 2*S*_*rh*_ into the exponential form gives

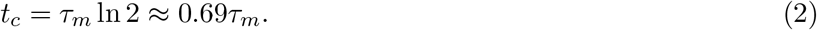

Peterchev et al. [16] estimated the strength-duration time constant for cortical magnetic stimulation at roughly 196 µs, with a 95% confidence interval of 181-210 µs, from motor threshold measurements obtained using controllable-pulse TMS. This value is influenced by multiple design choices made by the authors of the study and should not be interpreted as a universal neuronal membrane time constant.

### 2.1 Discharge circuit

We do not describe the precise wiring of the subcircuits in any of the following sections, as we assume the reader has access to a copy of the device’s schematic, which can be found at [tms_schematic.pdf](https://data.mendeley.com/datasets/gb8tgtfk3c/2/files/b9ef8c76-c0c9-4e88-886a-2dfddce42c6e). Additionally, we include a copy of the schematic in the supplementary materials. The following section describes the architecture and the reasons for each principal design choice. Building instructions are provided in Section 5.

The discharge circuit stores and releases the pulse energy. It is a series loop of the capacitor bank, the SCR module, and the TMS coil. We are interested in knowing how long it takes for the current to reach its peak value. This effective pulse length is obtained by analyzing the resulting underdamped RLC system when the capacitors are allowed to discharge freely through the coil. Since resistance in the underdamped discharge loop is not desired, we used the lossless LC system for our theoretical analyses as it provides a conservative upper bound on our circuit characteristics. The effective pulse lasts for a quarter of the period and its length is therefore given by

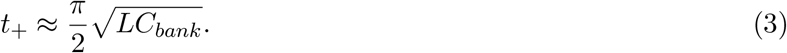

The coil is self-wound, so *L* varies between builds and must be measured (§5.2.5). We use a 230 µF capacitor bank and aim for a TMS coil with an inductance around 15 µH. This gives us an effective pulse length of around 90 µs.

The capacitor bank consists of two 460 µF polypropylene film capacitors wired in series, giving

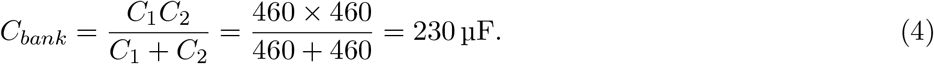

We use the KEMET C44UQGT6460T86K, a non-polarized DC link capacitor. Non-polarized parts are required for our proposed topology because the negative lobe can reach several hundred volts at maximum operating voltage. KEMET offers other parts in the C44U series at different ratings if the specific part we use is unavailable. The capacitor bank’s critical ratings are as follows:

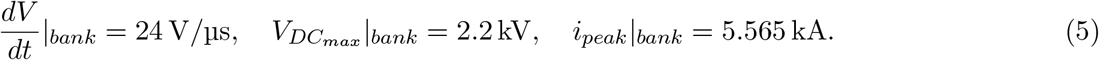

24 V/µs is obtained by summing the nominal 12 V/µs per-capacitor limit under equal voltage sharing, whereas the same series current flows through both capacitors and therefore the 5.565 kA is our specification of interest.

Let us define

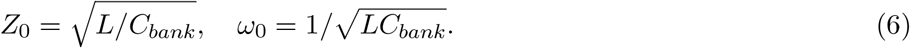

Using the characteristic quantities defined in Eq. (6), the lossless peak current which occurs at the quarter period or when the capacitor voltage crosses zero is given by Eq. (7).

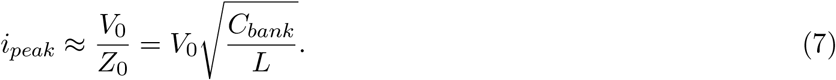

This is a conservative upper bound as the real peak will be slightly lower depending on the loop parasitic resistance.

The peak 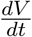 can be derived from the relation 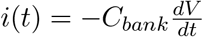,

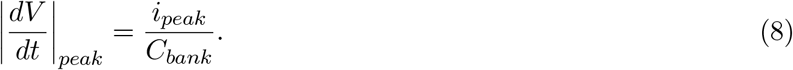

In the lossless system, the peak rate of voltage change coincides exactly with the peak current. Substituting the lossless *i*_*peak*_ from Eq. (7):

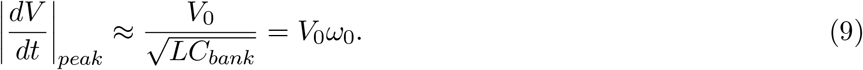

The 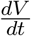 ceiling is therefore 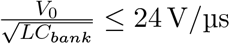, which sets the maximum operating voltage.

Assuming a TMS coil with *L* = 15.4 µH, we have *V*_0_ ≤ 1428.4 V for a perfectly lossless discharge system. This operating voltage also gives us *i*(*t*)_*peak*_ = 5520 A. Again, this is a conservative upper bound of the actual peak current as it assumes a lossless system. However, if deriving 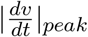 from measurements, a peak current of 5520 A will occur at 24 V/s even in a system with resistance. When operating this close to limits, be aware that you might be reducing your component’s lifetime and that catastrophic failures are more likely to happen. Always wear sufficient personal protective equipment and keep a physical barrier between you and the components. Recommended minimal personal protective equipment is described in section §5.1

To control the discharge circuit, we use the Littelfuse MCMA260PD1800YB, a SCR module integrating a thyristor and a diode. It can block up to 1800 V and handles forward surge currents of up to 8.97 kA. To survive this surge, the gate must receive at least 0.5 A at a 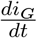 of at least 0.5 A/µs. These ratings drive the design of the gate driver (§5.3.4). The initial SCR current rise is: *V*_0_*/L* ≈ 97.4 A/µs at 1500 V. This is close to the SCR’s repetitive critical 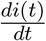 of 100 A/µs. The non-repetitive SCR 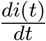 is 500 A/µs. These estimates also assume a strong gate signal. The module places the diode cathode at the thyristor anode node, which provides an antiparallel path required for the damping of the negative lobe.

Monophasic operation keeps only the positive lobe. The negative-lobe energy must be dissipated rather than returned to the bank. A damping resistor in series with the antiparallel diode carries the negative-lobe current and absorbs that energy, changing the system from an underdamped to an overdamped state once the capacitor voltage becomes negative. The resistor sees the full peak current, so it must have extremely high peak-power handling capacities. Since the required resistance is small (0.1 Ω), we fabricate the resistor from 1/8-inch wire rope. A good part of the capacitor bank energy is dissipated through this damping resistor. This means a significant amount of heat builds up in it. Since it is connected to the busbars, heat quickly spreads through the entire system. At the tested duty cycles, overheating was not an issue. Because thorough stress tests were not performed, we cannot recommend operations at rates higher than the tested duty cycles.

The TMS coil is a self-wound figure-of-eight coil with two wings, each with two stacked layers of five turns, similar in geometry to the MagVenture C-B60 coil. The self-made coil is inexpensive and allows testing of various designs and their resulting magnetic fields.

We place bleeder resistors across the capacitor terminals to form RC discharge paths with time constants much longer than that of the discharge loop, but short enough to bring the bank to a safe voltage in around 20 minutes if it is left charged.

Per-capacitor time constant:

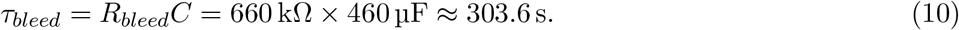

Assuming a *V*_0_ of 1500 V across the bank (750 V per capacitor), it takes around 17.2 min for the capacitor bank voltage to fall below 50 V.

Finally, a resistive voltage divider across the bank gives a low-voltage tap for the measurement and safety circuit (§5.4) and for general probing.

### 2.2 Control circuitry

The control circuitry provides isolated charge, fire, and trip signals from the host microcontroller to the discharge circuit. It sits in the power voltage domain and is driven from the control domain only through optocouplers, protecting the microcontroller and host computer.

The control circuitry consists of a relay and the following subcircuits built on separate perforated boards: an AND gate, linear voltage regulators, an optocoupler isolation board, and a gate driver.

A 24 VDC automotive relay (AZ9861-1C-24DC2R1) gates whether the PSU drives the charging circuit. The relay is normally open and its coil is controlled by an AND gate of two NPN BJTs so that the current flows only when `CHARGE CTRL` is high and `CMP STOP` is low. The `CMP STOP` signal is inverted as it passes through its respective optocoupler.

A 4700 µF electrolytic capacitor sits across the PSU terminals on the relay side, taped directly to the relay to minimize the inductance to the ZVS input. It supplies the high startup current the ZVS draws. An antiparallel Schottky diode across the relay coil suppresses the flyback voltage, protecting the driver circuitry.

The linear voltage regulators transform the PSU 24 V supply into PSU 12 V and PSU 5 V supplies, as labeled in the schematics. These regulators provide power to the rest of the control circuitry.

The optocoupler isolation board contains three optocouplers which transfer the CHARGE CTRL, FIRE CTRL, and `CMP STOP` signals.

The gate driver is built around the Microchip TC4420CPA MOSFET driver IC. The driver must turn the SCR on hard, which requires at least 0.5 A delivered at 0.5 A/µs into the gate. The TC4420 drives the primary of a 1:1 pulse transformer which carries the gate pulse to the high-voltage domain, driving the gate. Our gate driver provides a 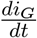 of 9.7 A/µs ≫ 0.5 A/µs (§5.3.5).

### 2.3 Measurement circuits

The measurement circuits contain two subcircuits that measure the capacitor bank voltage and share it with the user and the control circuitry. These circuits cross the high-voltage/control domain boundary. The high-voltage side of each is powered by a shared 9 V battery whose negative terminal is referenced to the negative capacitor terminal. This is done to maintain the floating-island topology. Communication to the control side occurs exclusively through optocouplers.

The comparator subcircuit prevents overcharging of the capacitor bank. An LM393 dual comparator monitors the bank voltage at the 1/401 (*V*_sense_) divider tap and disables charging once a configurable threshold is reached. The threshold trip acts on two independent paths. It drives a hardware shutoff through the `CMP STOP` signal into the relay AND gate, and it sets a software trip flag in the microcontroller. The hardware path stops charging with no software in the loop. The software flag adds a safety layer as it stops the user from sending the `CHARGE CTRL` signal and it logs the overvoltage. The trip threshold is set with a 5 kΩ potentiometer in a divider across the 9 V battery.

The linear optocoupler subcircuit transmits the voltage at the 1/401 (*V*_sense_) divider tap voltage from the high-voltage domain to the microcontroller without breaking isolation. It uses one LED and two photodiodes within a single linear-optocoupler package: a sender on the high-voltage side, a feedback receiver also on the high-voltage side to correct for the sender LED’s nonlinearity, and a receiver on the control side. This enables programmed charging up to a preset target voltage.

### 2.4 Charging circuit

The charging circuit raises the 24 V supply to the bank voltage. It is a three-stage chain: a ZVS driver, a self-wound step-up transformer, and a high-voltage rectifier. The step-up transformer can generate voltages over 1800 V from a bench supply at low cost.

We use a commercial two-inductor ZVS (Royer-type) driver. It self-oscillates into the transformer primary, so no external switching signal is needed. The two-inductor variant removes the center tap that a classic ZVS driver requires. With the primary connected, the driver resonates the 24 V input into a sine wave of peak

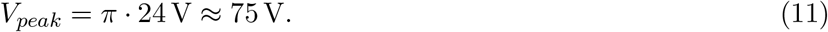

The sine wave has a frequency of around 30 kHz, making the step-up transformer smaller than it would be at line frequency. The output already exceeds safe touch voltage limits and is treated as hazardous. Operating the transformer near 30 kHz raises nerve- and cardiac-stimulation thresholds relative to 50 or 60 Hz, but does not confine contact current away from vital organs. IEC 60479-2:2019 [17] shows that for shock durations longer than the cardiac cycle, it takes approximately 14 times more current for cardiac fibrillation at 1000 Hz than at 50 − 60 Hz. The standard also shows that the ratio increases exponentially with frequency, although their analysis is limited to a maximal frequency of 1000 Hz.

The step-up transformer uses an ETD49 ferrite core with an air gap formed using a piece of paper approximately 0.1 mm thick. The gap lowers the inductance. The secondary is wound in five layers of 50 turns with a tap at the end of each layer (50, 100, 150, 200, 250 turns). Against the 10-turn primary, the taps give nominal step-up ratios of 5, 10, 15, 20, and 25, so at a 75 V primary peak, the tap peaks are approximately 375, 750, 1125, 1500, and 1875 V.

The operator selects the bank charge voltage by choosing a tap. This gives coarse voltage control. Finer control can then be achieved using the reading from the linear optocoupler measurement circuit.

The full-bridge rectifier converts the sine wave coming from the transformer secondary to DC. The DC output reaches the bank through current-limiting resistors. At the tested duty cycles, the resistors do not overheat.

### 2.5 Safety hardware and enclosure

The device operates from a few volts to over a thousand volts. **Inappropriate use is lethal**. Three safety layers act on the charging path. Two interlocks and a charge-cutoff switch sit in series with the PSU ahead of the relay: a lid interlock closed only when the enclosure covers the electronics, a dead-man interlock held by a second person, and a charge-cutoff switch. Any one of them opens the charging path. The comparator adds an overvoltage cutoff on both the hardware and software paths. These layers stop charging but do not drain the bank.

Two further safety layers drain the capacitor bank. The bleeder resistors discharge it passively over a few minutes, while a safety discharge stick gives a manual, rapid discharge option. A 1.5 m plastic pole carries an insulated lead to a hooked bare end and returns through two 1 kΩ 50 W resistors permanently tied to the bank’s negative terminal. The stick lets the operator discharge the bank through a hole in the enclosure wall without having to open it and while being able to keep a safe distance from the device.

The enclosure is a transparent box, so the operator can watch for sparks or motion while the device is energized. The discharge circuit sits on a Styrofoam platform that isolates it from the box and the bench. At maximum voltage, the Lorentz forces between opposing conductors are large, so junctions such as the coil-lead Y-split are mechanically restrained. Optionally, a lid-mounted fan handles cooling. Heat was not a limiting factor at the pulse rates tested.

### 2.6 Software

An Arduino UNO R4 Minima microcontroller communicates with an application on the operator’s computer over a single USB serial link at 115200 baud using newline-terminated ASCII messages. Capacitor voltage thresholds can be set in the microcontroller firmware, which will automatically shut down charging if it detects a capacitor charge higher than the preset maximum. The host issues commands to manually activate or deactivate charging, arm the device, and fire the pulse. The full command and message sets are listed in §6.2 (Tables 4 and 5).

Additionally, we provide a data acquisition script which interfaces with an oscilloscope over a serial link to make the capture of the full 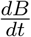 maps easier.

Finally, a Jupyter Notebook contains the step-by-step instructions accompanied by code to characterize the device based on the collected field maps. The data analysis outputs an estimated TMS coil 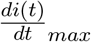, which we then feed into SimNIBS to estimate peak *E*-fields on a default anatomy.

## 3 Design files summary

All design files and accompanying data are available in the [online repository](https://data.mendeley.com/datasets/gb8tgtfk3c/2).

### KiCAD schematic files

**discharge.kicad sch** - Schematic overview of the entire TMS circuit with a focus on the TMS discharge circuit.

**charge.kicad sch** - Main charging circuit schematic with measurements and control sub-circuits.

**gate driver.kicad sch** - Gate driver sub-circuit schematic.

**tms schematic.pdf** - Entire TMS circuit schematic in PDF format.

### LTSpice schematic files

**discharge.asc** - TMS inductor-capacitor oscillator discharge circuit schematic for SPICE simulation.

**charge.asc** - ZVS capacitor charging circuit schematic for SPICE simulation.

**gate driver.asc** - Gate driver circuit schematic for SPICE simulation.

**linear optocoupler.asc** - Linear optocoupler measurement circuit schematic for SPICE simulation.

### FreeCAD parametric CAD files

**coil winder.FCStd** - Coil winder helper for winding the TMS coil.

**pickup cylinder.FCStd** - Small cylinder to help wind the pickup coil.

**sensing cross.FCStd** - Sensing cross to translate user movements to pickup coil movements.

**busbars.FCStd** - 2D busbar drawings with dimensions.

**coil windings.FCStd** - 2D representation of TMS coil windings.

**TMS field grid.pdf** - Measurement grid for optimal pickup coil positioning.

### Safety checklists

**Safety Checklists.pdf** - Device inspection, pre-charge, and de-energization safety checklists. Also available in the supplementary materials.

### Control software files

**controller arduino.ino** - Arduino sketch for controlling the TMS circuit.

**app.py** - Python script for the user interface.

### Data collection and analysis files

**collect grid.py** - Software interface with the oscilloscope for collecting pick up coil measurements.

**data analysis.ipynb** - Jupyter notebook for analyzing the data acquired with the TMS.

**create fo8 coil.py** - Code modified from SimNIBS to generate a figure-of-eight coil model.

## 4 Bill of materials summary

The bill of materials (BOM) reflects component prices as of June 2026. It is separated into five parts. The first three reflect the three different pages of the schematics. The last two are Sensing and Miscellaneous, respectively. Sensing contains components required in §7 Validation, whereas Miscellaneous contains components that could not fit neatly into one of the preceding parts of the table.

The bill of materials includes the components and materials incorporated into the device, but excludes reusable laboratory instruments, fabrication tools, computing resources, and institutional safety infrastructure. Access to the items in Table 3 is therefore assumed.

**Table 1:**
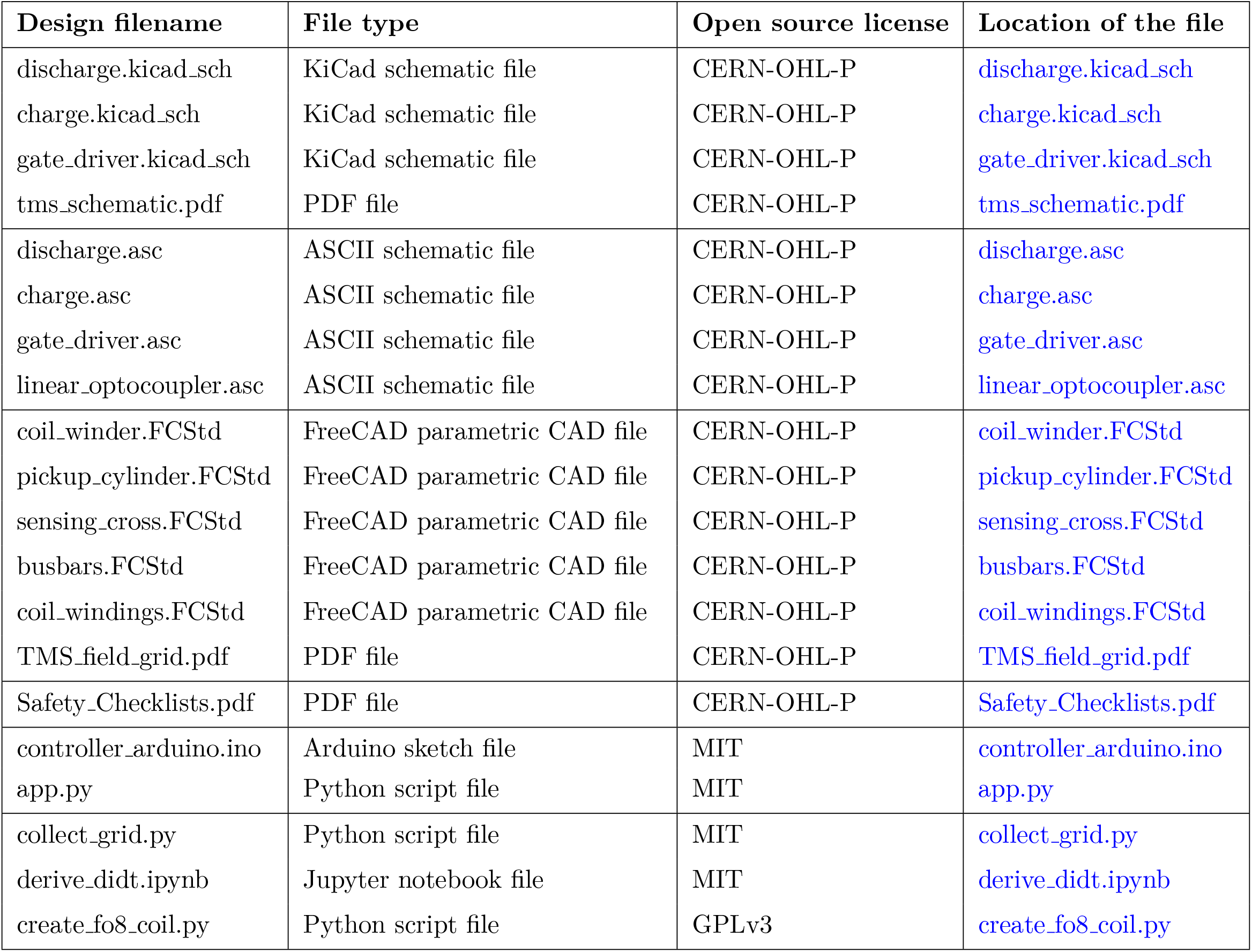
Summary of the design files provided in the project repository.

**Table 2:** Bill of materials.

| Designator | Component | Qty | Cost per unit (USD) | Total cost (USD) | Source of materials | Material type |
| --- | --- | --- | --- | --- | --- | --- |
| **Discharge circuit** |  |  |  |  |  |  |
| C1, C2 | C44UQGT6460T86K<br>Main discharge capacitors 460 uF each | 2 | 110.56 | 221.12 | DigiKey | Metal |
| R1, R2, R3, R4, R5, R6 | FMP200JT-52-220K<br>Bleed resistors 220k Ohm 2W | 6 | 0.20 | 1.20 | DigiKey | Metal |
| D1,Q1 | MCMA260PD1800YB<br>SCR Module | 1 | 119.78 | 119.78 | DigiKey | Composite |
| R7 | Damping resistor<br>1/8" (3MM) Wire Rope, 100FT<br>Stainless Steel Cable | 1 | 16.99 | 16.99 | Amazon | Metal |
|  | Busbar wiring<br>Copper Pipe Type M 3/4-inch x 5 Foot Straight Length | 1 | 26.81 | 26.81 | Home Depot | Metal |
|  | Busbar isolation<br>Kapton tape | 1 | 2.50 | 2.50 | Amazon | Polymer |
| R18 | 1M Ohm 1/4W resistor<br>Voltage divider high side resistors | 10 | 0.10 | 1.00 | DigiKey | Ceramic |
| R19 | 22k Ohm 1/4 resistor<br>Voltage divider low side resistor | 1 | 0.10 | 0.10 | DigiKey | Ceramic |
| R19 | 3k Ohm 1/4 resistor<br>Voltage divider low side resistor | 1 | 0.10 | 0.10 | DigiKey | Ceramic |
| L1 | Coil wire<br>10 AWG Stranded copper wire (25ft) | 1 | 14.99 | 14.99 | Amazon | Metal |
|  | Wire insulation<br>Kapton tape | 1 | 2.50 | 2.50 | Amazon | Polymer |
|  | Coil insulation<br>Electrical tape | 1 | 3.78 | 3.78 | Amazon | Polymer |
|  | Coil winder filament<br>85g of PLA filament. 1kg for 20.39 cad. 1kg for 13.99 usd. | 1 | 1.19 | 1.19 | Amazon | Polymer |
| SW1 | On/Off Switch<br>DC On/Off switch for PSU | 1 | 14.44 | 14.44 | DigiKey | Composite |
| SW2, SW3 | Contact switches (microwave door switches)<br>Interlock switches | 2 | 2.97 | 5.94 | Amazon | Composite |
| R8, R9 | 50W 1k Ohm discharge resistor<br>Safety stick discharge resistors | 2 | 5.99 | 11.98 | DigiKey | Ceramic |
|  | PVC pipe<br>PEX-B pipe for safety stick | 1 | 3.22 | 3.22 | Home Depot | Polymer |
|  | 18 AWG HV silicone wire<br>Safety stick cable (5ft) | 5 | 1.41 | 7.05 | DigiKey | Polymer |
| **Gate driver circuit** |  |  |  |  |  |  |
| U12 | TC4420CPA<br>MOSFET gate driver | 1 | 2.02 | 2.02 | DigiKey | Composite |
|  | A 08-LC-TT<br>8 pin IC socket (optional) | 5 | 0.67 | 3.35 | DigiKey | Composite |
| C14 | 0.1 uF ceramic capacitor<br>Fast energy bank for the pulse | 1 | 0.24 | 0.24 | DigiKey | Ceramic |
| C15 | 100 uF electrolytic capacitor<br>Slow energy bank for the pulse | 1 | 0.53 | 0.53 | DigiKey | Metal |
| T2 | P0584NL<br>1:1 Pulse Transformer | 1 | 4.77 | 4.77 | DigiKey | Composite |
| D19, D20, D21, D22 | 1n5819<br>Schottky Diodes | 4 | 0.29 | 1.16 | DigiKey | Semiconductor |
| R30 | 10 Ohm gate resistor<br>10 Ohm gate resistor | 1 | 0.10 | 0.10 | DigiKey | Ceramic |
|  | 22 AWG wire<br>Wire for the whole circuit (1 m) | 1 | 0.30 | 0.30 | Amazon | Metal |
| U11 | VO2611<br>HV Pulse Optocoupler | 1 | 2.25 | 2.25 | DigiKey | Composite |
| R28 | 620 Ohm 1/4W resistor<br>620 Ohm resistor for optocoupler LV side | 1 | 0.10 | 0.10 | DigiKey | Ceramic |
| R29 | 1.2k Ohm 1/4W resistor<br>1.2k Ohm resistor for optocoupler HV side | 1 | 0.10 | 0.10 | DigiKey | Ceramic |
| C13 | 0.1 uF optocoupler capacitor<br>0.1 uF capacitor for optocoupler HV side | 1 | 0.24 | 0.24 | DigiKey | Ceramic |
| **Charging circuit** |  |  |  |  |  |  |
| U1 | L7812CV<br>Linear voltage regulator | 1 | 1.17 | 1.17 | DigiKey | Composite |
| U2 | L7805CV<br>Linear voltage regulator | 1 | 1.05 | 1.05 | DigiKey | Composite |
| C5, C6 | 0.33 uF ceramic capacitor<br>Supports the voltage regulators | 2 | 0.32 | 0.64 | DigiKey | Ceramic |
| C7, C8 | 0.1 uF ceramic capacitor<br>Supports the voltage regulators | 2 | 0.24 | 0.48 | DigiKey | Ceramic |
| A1 | Arduino UNO R4 Minima<br>Arduino | 1 | 20.00 | 20.00 | DigiKey | Composite |
| K1 | AZ9861-1C-24DC2R1<br>24 VDC relay | 1 | 5.66 | 5.66 | DigiKey | Composite |
| D2 | 1n5819<br>Schottky Diode | 1 | 0.29 | 0.29 | DigiKey | Semiconductor |
| C9 | 1189-1397-ND<br>4700 uF 50V capacitor | 1 | 3.50 | 3.50 | DigiKey | Metal |
| ZVS Driver | ZVS Driver<br>ZVS Driver | 1 | 15.99 | 15.99 | Amazon | Composite |
| T1 | ETD49/25/16-3C94<br>Transformer Cores | 2 | 3.93 | 7.86 | DigiKey | Ceramic |
| T1 | B66368W1020T001<br>Transformer Bobbin | 1 | 4.14 | 4.14 | DigiKey | Polymer |
| T1 | B66368A2000X000<br>Transformer Clips | 2 | 0.80 | 1.60 | DigiKey | Metal |
| D3, D4, D5, D6, D7, D8, D9, D10, D11, D12, D13, D14, D15, D16, D17, D18 | UF4007<br>HV Diode | 16 | 0.50 | 8.00 | DigiKey | Semiconductor |
| R16, R17 | 1k Ohm 25W<br>Resistor | 2 | 5.22 | 10.44 | DigiKey | Ceramic |
| U8A | LM393N/NOPB<br>Voltage Comparator | 1 | 0.92 | 0.92 | DigiKey | Composite |
| RV1 | 5k Ohm potentiometer<br>Comparator voltage divider<br>potentiometer | 1 | 5.35 | 5.35 | DigiKey | Composite |
| R23 | 5.6k Ohm resistor<br>Comparator voltage divider<br>resistor | 1 | 0.10 | 0.10 | DigiKey | Ceramic |
| R24 | 1M Ohm resistor<br>Hysteresis resistor | 1 | 0.10 | 0.10 | DigiKey | Ceramic |
| R25 | 10k Ohm resistor<br>Comparator pull up resistor | 1 | 0.10 | 0.10 | DigiKey | Ceramic |
| BT1 | 9V battery<br>Comparator battery | 1 | 1.59 | 1.59 | Amazon | Metal |
| U3, U4, U9 | VO2611<br>HV Pulse Optocoupler | 3 | 2.25 | 6.75 | DigiKey | Composite |
| R10, R13, R27 | 620 Ohm 1/4W resistor<br>620 Ohm resistor for optocoupler<br>LV side | 3 | 0.10 | 0.30 | DigiKey | Ceramic |
| R11 | 1.2k Ohm 1/4W resistor<br>1.2k Ohm resistor for optocoupler<br>HV side | 1 | 0.10 | 0.10 | DigiKey | Ceramic |
| R26 | 1.2k Ohm 1/4W resistor<br>1.2k Ohm resistor for optocoupler<br>LV side | 1 | 0.10 | 0.10 | DigiKey | Ceramic |
| C10, C12, C14 | 0.1 uF optocoupler capacitor<br>0.1 uF capacitor for optocoupler<br>HV side | 3 | 0.24 | 0.72 | DigiKey | Ceramic |
| C11, C13, C15 | 22 pF optocoupler capacitor<br>22 pF capacitor for optocoupler<br>HV side | 3 | 0.30 | 0.90 | DigiKey | Ceramic |
| Q2, Q3 | BC337 NPN Transistor<br>NPN for relay coil control | 2 | 0.18 | 0.36 | DigiKey | Semiconductor |
| R12, R14 | 1.5k Ohm 1/4W resistor<br>Base resistor for NPN | 2 | 0.10 | 0.20 | DigiKey | Ceramic |
| R3 | 10k Ohm 1/4W resistor<br>Pull down NPN resistor | 1 | 0.10 | 0.10 | DigiKey | Ceramic |
|  | 16 AWG wire<br>PSU to ZVS wiring | 1 | 3.33 | 3.33 | Amazon | Metal |
|  | 28 AWG enameled wire<br>Primary and secondary wiring (530 ft) | 1 | 11.99 | 11.99 | Amazon | Metal |
|  | 18 AWG HV silicone wire<br>Full bridge to capacitor wiring (1ft) | 1 | 1.41 | 1.41 | DigiKey | Polymer |
|  | 22 AWG wire<br>Wire for LV circuit boards (1 m) | 1 | 0.30 | 0.30 | Amazon | Metal |
| U6 | IL300<br>Linear Optocoupler | 1 | 3.81 | 3.81 | DigiKey | Composite |
| U5, U7, U1 | UA741<br>Op amp for linear optocoupler | 3 | 0.64 | 1.92 | DigiKey | Composite |
| R20, R22 | 100k Ohm resistor<br>100k Ohm gain resistors for linear optocoupler | 2 | 0.10 | 0.20 | DigiKey | Ceramic |
| R21 | 1k Ohm resistor<br>1k Ohm servo resistor | 1 | 0.10 | 0.10 | DigiKey | Ceramic |
| U8A | LM358<br>Op amp for voltage follower | 2 | 0.30 | 0.60 | DigiKey | Composite |
|  | Dupont wires MM<br>Wires to Arduino | 20 | 0.06 | 1.20 | Amazon | Metal |
| **Sensing** |  |  |  |  |  |  |
|  | 28 AWG wire<br>Pickup coil (2ft), included in the transformer winding wire | 1 | 0.50 | 0.50 | Amazon | Metal |
|  | PLA filament<br>PLA for winding cylinder (2g) | 1 | 0.05 | 0.05 | Amazon | Polymer |
|  | 100x P4250 Probe<br>100x oscilloscope probe with 2000Vpp rating | 1 | 22.99 | 22.99 | Amazon | Composite |
|  | PLA filament<br>PLA for sensing cross (6g) | 1 | 0.15 | 0.15 | Amazon | Polymer |
| **Miscellaneous** |  |  |  |  |  |  |
|  | Perforated boards | 6 | 0.25 | 1.50 | Amazon | Composite |
|  | M6 bolts | 2 | 2.97 | 5.94 | Home Depot | Metal |
|  | M6 nuts | 2 | 3.97 | 7.94 | Home Depot | Metal |
|  | M6 washers<br>Large washers | 2 | 0.21 | 0.42 | Home Depot | Metal |
|  | Enclosure<br>50L container | 1 | 20.98 | 20.98 | Home Depot | Polymer |
|  | Zip ties | 1 | 3.99 | 3.99 | Amazon | Polymer |
| **Total (USD, before tax)** |  |  |  | **657.68** |  |  |

**Table 3:** Tools, equipment, and infrastructure not included in the bill of materials.

| Category | Item | Use |
| --- | --- | --- |
| Electrical instrumentation | Adjustable 24 V DC bench power supply | Must provide adjustable current limiting for low-voltage checkpoints and tolerate the approximately 10 A ZVS startup transient. |
| Electrical instrumentation | Digital storage oscilloscope | At least two input channels and single-shot acquisition for measuring the capacitor-discharge and gate-driver waveforms. USB/SCPI communication is required to use the supplied automated field-mapping script without modification. |
| Electrical instrumentation | Standard oscilloscope probes | At least two probes suitable for low-voltage circuit measurements. |
| Electrical instrumentation | Digital multimeter and test leads | Required for resistance, continuity, and voltage measurements. |
| Electrical instrumentation | Function generator | Must produce an adjustable DC level below 5 V and a 0–5 V, 1 kHz sawtooth waveform for Checkpoint C. |
| Electrical instrumentation | LCR meter | Optional. Used to measure the self-wound coil inductance and resistance. Inductance can alternatively be estimated from the measured discharge waveform. |
| Fabrication | 3D printer or printing service | Required to fabricate the TMS coil winder, pickup-coil former, and pickup-coil positioning guide. The cost of plastic is included in the BOM. |
| Fabrication | Soldering station | Required for circuit-board assembly. |
| Fabrication | Solder, flux, and fume extraction | Consumable soldering supplies and appropriate ventilation. |
| Fabrication | Electric drill and drill bits | Includes the 1/16-inch and 1/4-inch bits used for the busbars and suitable tools for making openings in the enclosure. |
| Fabrication | Hacksaw | Used to cut the copper plumbing pipe into busbar sections. |
| Fabrication | Pliers | Used to flatten the copper pipe, bend busbars, manipulate conductors, and wind the coil. |
| Fabrication | Screwdrivers and wrenches | Required for the M6 capacitor and SCR busbar connections. |
| Fabrication | Emery cloth, file, or deburring tool | Used to remove coatings and oxidation, expose enameled wire, and deburr drilled or cut metal. |
| Personal Protective Equipment | Class 1 rubber insulating gloves compliant with ASTM D120 or IEC 60903 | Gives some protection against the voltages used in the build. |
| Personal Protective Equipment | Impact-rated eye protection | Gives some protection against potential debris ejection. |

## 5 Build instructions

The TMS contains multiple subsystems that can and should be independently built and tested. Figure 3 contains a high-level system diagram showing the different parts of the build and their respective voltage domains/isolation layers. The recommended build order is:

**Figure 3:**
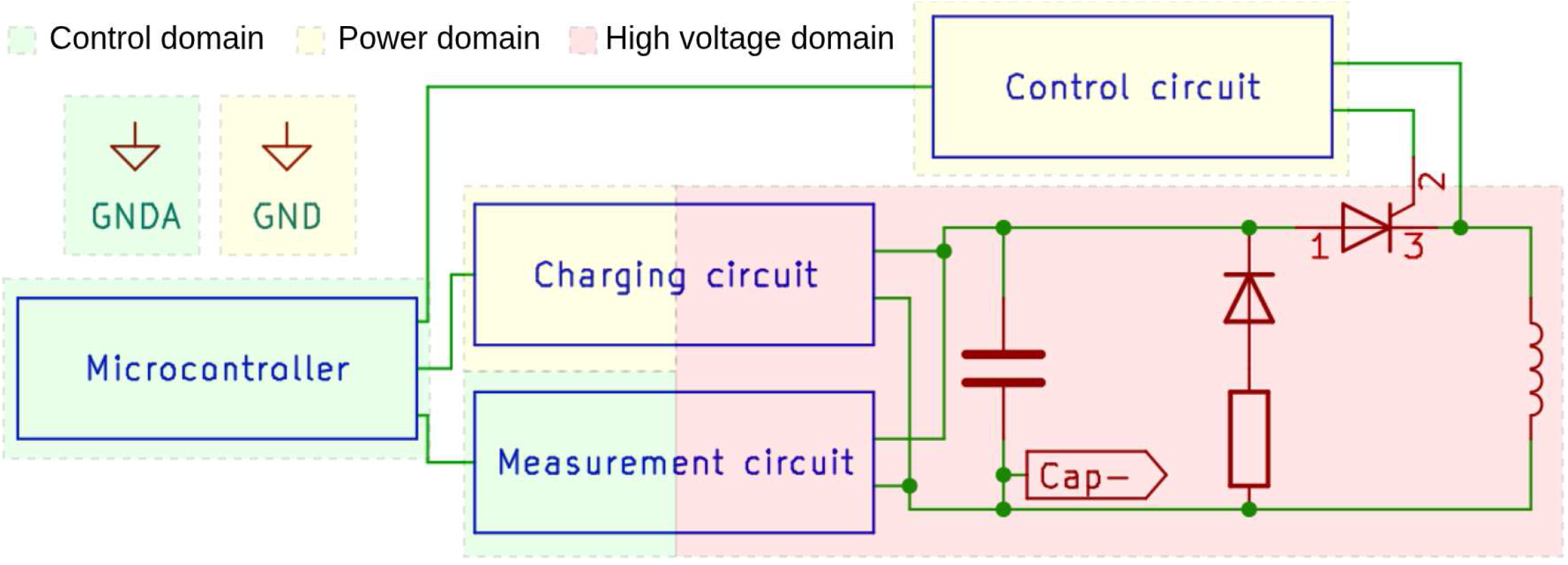
Systems overview. Block diagram of the different subsystems and their voltage domains. The control domain is highlighted in green, the power domain is highlighted in yellow, and the high-voltage domain is highlighted in red. The control domain contains the microcontroller and a part of the measurement circuits which is isolated from the rest by optocouplers. The power domain contains the PSU, control circuitry, and a part of the charging circuit. The microcontroller communicates with the charging and control circuits via optocouplers. The charging and control circuits are galvanically isolated from the high-voltage domain since they use transformers. In the schematics, the `GNDA` symbol represents the control domain common ground, the `GND` symbol represents the power domain common ground, and CAP-represents the high-voltage-domain common ground.

1. Discharge circuit (§5.2)
2. Control circuitry (§5.3)
3. Measurement circuits (§5.4)
4. Charging circuit (§5.5)
5. Safety hardware and enclosure (§5.6)

### 5.1 Safety concerns

The hardware described here operates from a few volts to over a thousand volts. It contains a high-energy capacitor bank, generates high voltages, and creates strong, rapidly changing magnetic fields. **Inappropriate use is lethal**. Contact with the system can cause severe injury or death, and stored energy can remain after charging is halted. Never rely on the computer interface or firmware as the primary safety mechanism. Work must be performed by trained personnel. Do not connect this research prototype to a person or animal. Never operate the device alone or without appropriate protective equipment when lethal voltages are present. Personal protective equipment (PPE) is the last line of defense, not a substitute for proper use of the safety mechanisms and remote operation. The design includes multiple, low-cost safety layers; do not skip any of them.

Whenever handling high voltages, we recommend using Class 1 rubber insulating gloves compliant with ASTM D120 or IEC 60903, insulated footwear, and that everyone present wears impact-rated eye protection.

The instructions below describe how to build each subsystem and validate it at low voltages before moving on. Do not energize the charging circuit at high voltage until every subsystem has passed its low-voltage checkpoint and all safety hardware is installed.

Additionally, we provide a safety checklist to be completed before every energization of the high-voltage system, and a checklist for each de-energization ([Safety_Checklists.pdf](https://data.mendeley.com/datasets/gb8tgtfk3c/2/files/c0abcd49-44b4-44b4-abee-3f267c929052)).

### 5.2 Discharge circuit

The discharge circuit is an RLC oscillator consisting of the capacitor bank in series with the SCR module and TMS coil. The bank capacitance and the coil inductance set the pulse length and the required component ratings (design choices explained in §2.1). Figure 4 shows the isolated discharge circuit.

**Figure 4:**
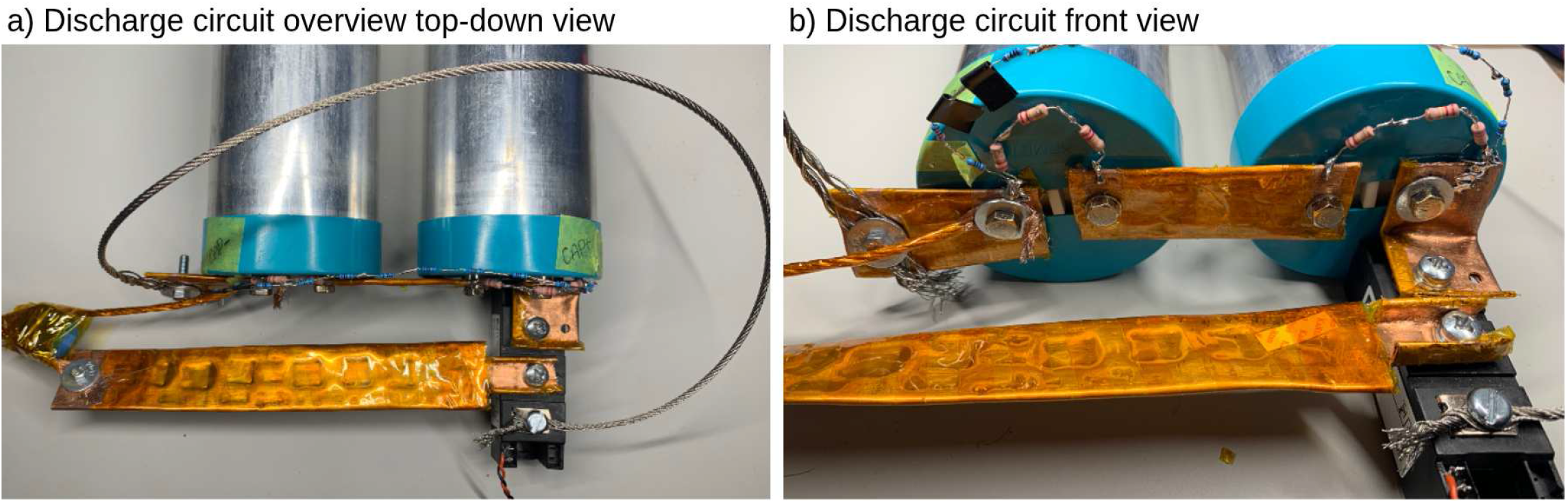
Overview of the discharge circuit. **a) Discharge circuit overview top-down view**. Two capacitors are connected in series with an SCR module and the TMS coil. A damping resistor is placed in series with a diode antiparallel to the discharge LC circuit. **b) Discharge circuit overview front view**. Bleeder resistors are placed across each capacitor’s terminals to drain them over a few minutes. A voltage divider consisting of 12 resistors in series is connected across the terminals of the capacitor bank.

#### 5.2.1 Busbars

The four busbars carrying high pulse currents are improvised from type-M 3/4-inch copper plumbing pipe to keep costs low. Pipe is cut to length with a hacksaw and flattened with pliers. Be careful not to overflatten the edges as cracks in a busbar disqualify it from use. Protective and oxidation coatings are removed with emery cloth at connection sites. Mounting holes are drilled in stages from a 1/16-inch pilot up to the final 1/4-inch diameter that accepts the M6 nuts supplied with our capacitors and SCR module. Busbar dimensions are presented in Figure 5.

**Figure 5:**
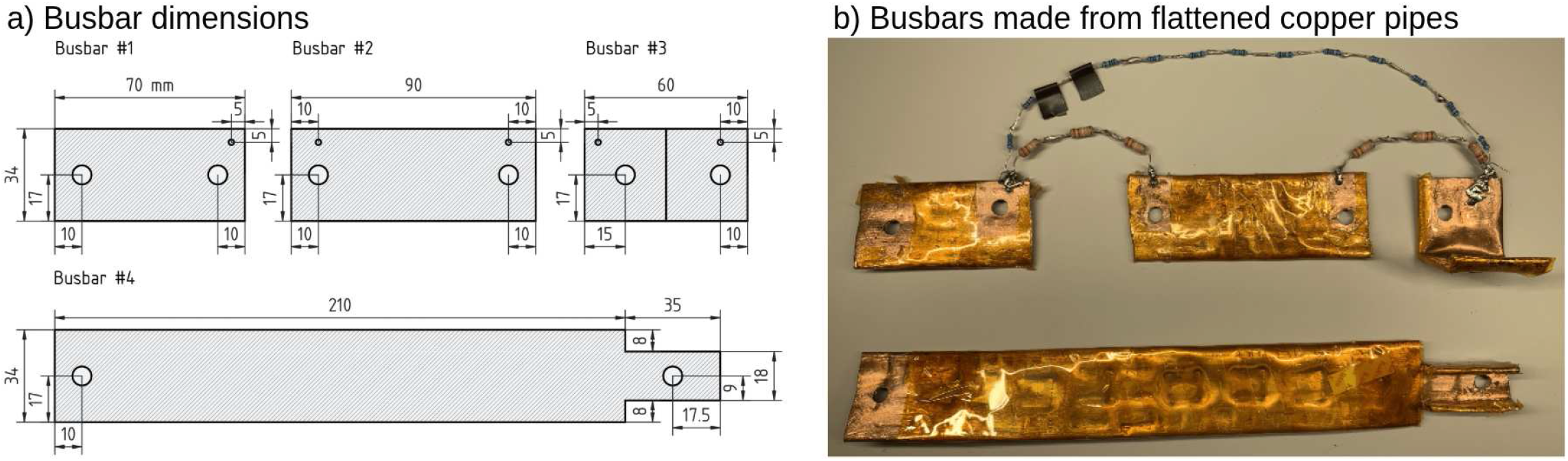
Busbar construction. **a) Busbar dimensions**. 2D drawing with dimensions of the four busbars. Dimensions are in millimeters. Busbar #3 has a 90° bend in the middle. **b) Busbars made from flattened copper pipes**. The busbars used in the discharge circuit were created by flattening copper pipes and cutting them to the prescribed dimensions. The bleeder resistors—a series connection of three 220 kΩ resistors—and the voltage divider are installed using the 1/16-inch corner holes.

The Lorentz forces produced during firing further flatten the busbars. Although this deformation is mechanically acceptable, extended use could crack the pipe along the flattened edges. If cracking is noticeable, parasitic resistance rises and power delivered to the coil drops. Regularly inspect the busbars after firing tests. We did not encounter issues with this setup during our tests. All busbars are wrapped in two layers of Kapton tape except at connection sites.

The four busbars and their roles:

1. **Busbar #1** serves as a hub on the capacitor bank negative terminal.
2. **Busbar #2** serves as a series link between the two capacitors.
3. **Busbar #3** connects the capacitor bank positive terminal to the SCR anode. It contains a 90° bend at its midpoint.
4. **Busbar #4** connects the SCR cathode to one coil lead; one edge is narrowed to maintain creepage from busbar #3.

On #1, #2, and #3, we also drill 1/16-inch auxiliary holes near a top corner to simplify connecting the bleeder and voltage-divider wires. A second auxiliary hole is drilled on the lower half of busbar #3 to allow for better contact when discharging the capacitor bank with the safety stick.

#### 5.2.2 Damping resistor

Using a multimeter in the appropriate resistance range, measure the resistance of a 10 m length of your wire rope to obtain its per-meter resistance, then cut a length giving the target 0.1 Ω. For our batch, we needed 76 cm. Sand the individual strands at the cut ends with emery cloth to remove any coating before making connections. Screws are inserted and secured between the wire rope strands.

#### 5.2.3 Capacitor bank

The two capacitors are connected in series using busbar #2. Busbar #1 is connected to the left-most capacitor terminal (bank negative). Busbar #3 is connected to the right-most capacitor terminal (bank positive); its 90° bend sits below the terminal with the lip parallel to the bench.

#### 5.2.4 SCR module

Mount the Littelfuse MCMA260PD1800YB SCR module on the platform beside the capacitor bank. Bolt busbar #3 to the module anode and the narrow end of busbar #4 to the module cathode with the supplied M6 bolts. Keep the gate and auxiliary-cathode terminals clear and accessible. Confirm the module pinout against the datasheet before bolting. If the module you use has a different pinout, you might need to change the busbar dimensions and physical layout.

#### 5.2.5 Coil construction

The coil described in this section is a low-cost prototype intended to demonstrate the operation of the stimulator. It is not presented as a durable, clinically qualified, or human-use stimulation coil. We expect researchers using this system to design and validate coils appropriate to their own applications. The manually applied polyimide- and electrical-tape insulation has a limited and uncharacterized service life. Kapton tape can become brittle with age; handling, thermal cycling, and mechanical stresses may also cause cracking of the insulation, changes of the coil shape, or abrasion of the insulating layers. Inspect the complete coil, lead junctions, and insulation layers before every use. Do not energize the coil if any deterioration, exposed conductors, unusual deformation, or arcing is observed. We did not perform accelerated aging tests. The coil should be treated as a consumable prototype and be rebuilt whenever deterioration is observed or before reuse following prolonged storage.

Strip 7.62 m (25 ft) of 10 AWG (2.59 mm diameter) stranded copper wire down to bare copper, then reinsulate the full length with Kapton tape with 50% overlap, giving two layers everywhere. A 3D-printed coil winder ([coil_winder.FCStd](https://data.mendeley.com/datasets/gb8tgtfk3c/2/files/5ede680a-900a-4428-a39e-35d5ee84654a)) helps you follow the geometry during winding. This coil is a benchtop prototype and is not suitable for placement on a person.

Leave approximately 1 m of wire as a lead, then wind the two wings as shown in Figure 6. Starting at the bottom layer of the right wing, wind five turns from the outside inward before going up to the top layer of the right wing and winding five turns from the inside outward. Use short strips of electrical tape to hold turns in place as you go. The two wings must be wound so that the wires passing through the central focal region all carry current in the same direction. Carry the wire from the top of the top layer of the right wing to the bottom of the bottom layer of the left wing. From there, wind five turns from the outside inward on the bottom layer before winding from the inside outward on the top layer. Finally, carry the wire down through the hole at the bottom of the coil winder until the wire meets the opposite lead. Twist the two leads tightly together immediately as they exit the coil to reduce stray inductance, then add two layers of Kapton tape over the twisted section to hold it and further isolate it.

**Figure 6:**
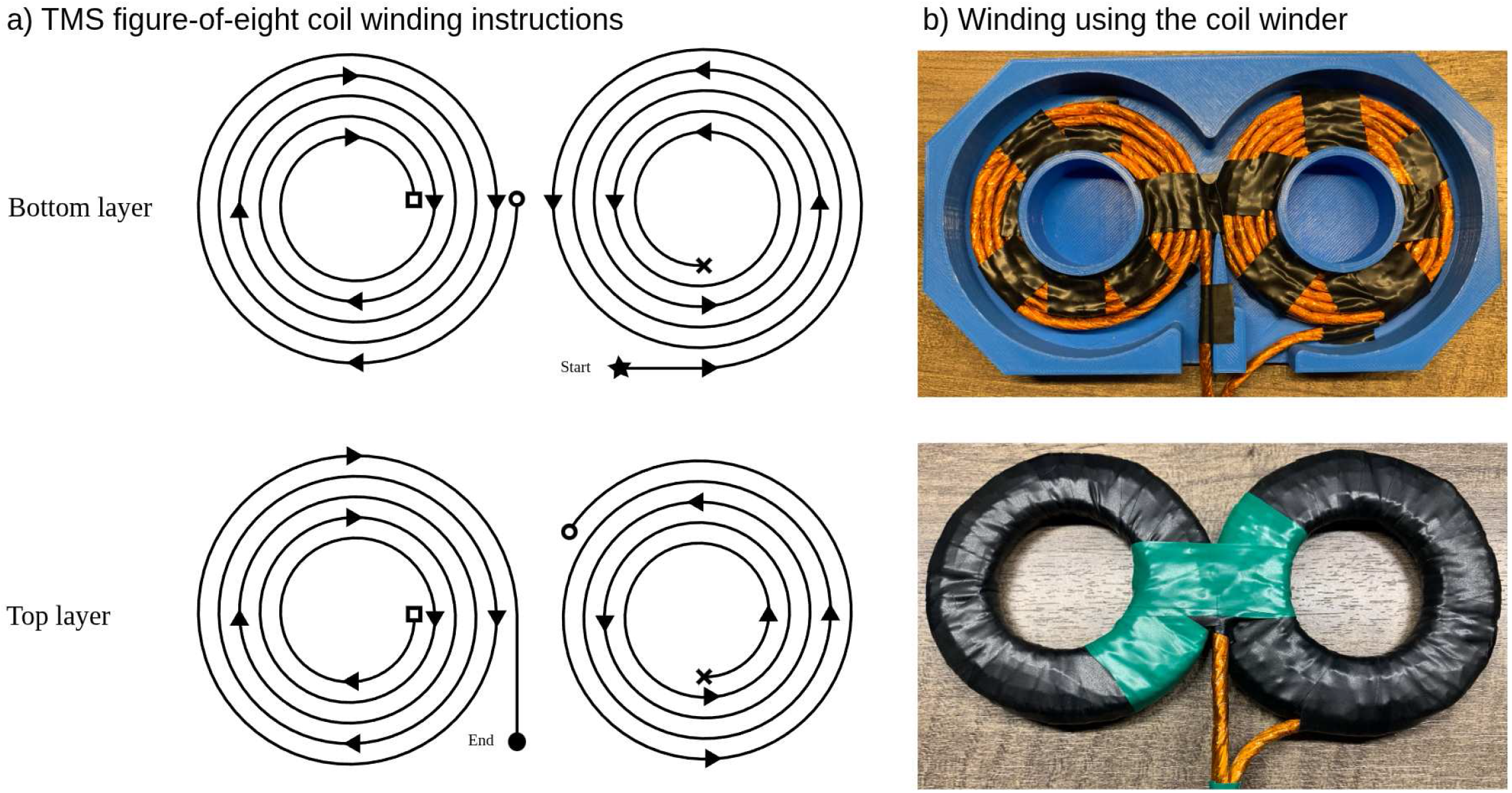
Winding the TMS figure-of-eight coil. **a) TMS figure-of-eight coil winding instructions**. Top-down view of the coil’s windings. Starting at the bottom layer on the right wing, the wire creates five turns before transitioning to the top layer as indicated by the cross mark. Similarly, the wire later transitions from the top to the bottom layer indicated by the circle symbols. The last transition from the bottom to the top layer is indicated by the square symbols. **b) Winding using the coil winder**. Images of an actual coil wound following the winding instructions with the help of our coil winder object ([coil_winder.FCStd](https://data.mendeley.com/datasets/gb8tgtfk3c/2/files/5ede680a-900a-4428-a39e-35d5ee84654a)).

Opposing currents in the leads produce a Lorentz force that tries to separate them. In practice, we found that the two layers of Kapton tape withstood the forces without visible deformation. Movement in the coil was imperceptible. Carefully remove the coil from the winder and use electrical tape across the wings to improve the structural integrity of the coil and remove residual gaps in the turns. The coil stores a large amount of magnetic energy, and a coil failure can eject conductors at high speeds. Always wear eye protection and further enclose the coil for initial tests.

Because the coil is self-wound, inductance varies between builds. Measure *L* with an LCR meter and adjust your maximum operating voltage so as not to exceed the bank 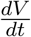 rating. If you do not have access to an LCR meter, you can estimate *L* during Checkpoint A §5.2.8 by using the quarter lobe duration and the known capacitance. Always test your coil at low voltages first and step up gradually. If you observe sparks or visible motion, stop, reinforce the insulation, or rebuild the coil.

Our finalized coil is 14.9 cm wide, 7.5 cm high, and 0.8 cm deep. Measured inductance: 15.4 µH. Measured resistance at 3 kHz: 30.4 mΩ. We chose 3 kHz because the underdamped discharge frequency is approximately 2710 Hz with a 230 µF capacitor bank and a 15 µH inductor.

#### 5.2.6 Bleeder resistors

Each capacitor receives three 220 kΩ 2 W resistors in series, giving 660 kΩ per capacitor; six resistors total. **Do not substitute** a single 660 kΩ resistor for the three-resistor string. The working voltage across one capacitor at full charge exceeds the working voltage rating of most single 2 W resistors. At the capacitor bank’s maximal rated voltage of 2.2 kV, which exceeds the downstream circuit limits but provides an upper bound for alternative design, each resistor dissipates 0.61 W.

Solder three resistors in series to form one bleeder string, build two, and connect each across the terminals of the capacitors. Sand the busbars at the solder sites; the high thermal mass of the busbar will require significant heating for solder to flow. Ensure the connection is solid.

#### 5.2.7 Voltage divider

The high side of the divider consists of ten 1 MΩ 1/4 W resistors in series. The low side consists of a 22 kΩ and a 3 kΩ 1/4 W resistor in series. The low side is connected to CAP-, while the high side is connected to CAP+. **Do not substitute** the ten 1 MΩ resistors with fewer, higher-value parts. 1/4 W resistors typically have low working-voltage ratings.

Two taps are taken:

- *V*_**sense**_ between the high and low sides for the comparator and linear optocoupler circuits. It provides a 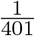 ratio of the capacitor voltage.
- *V*_**1M**_ after the first 1 MΩ resistor above CAP-for multimeter probing. A *V*_**2M**_ tap, located two 1 MΩ resistors above CAP-, is used when testing at low voltages.

Without an attached meter, the two monitoring-tap ratios are

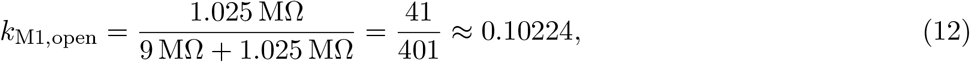

and

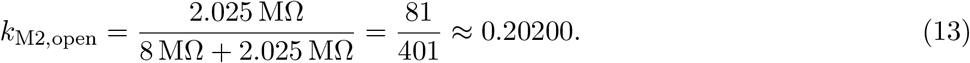

These are approximately 1/10 and 1/5. Connecting a multimeter between either tap and CAP-places the meter input resistance in parallel with the divider resistance below that tap. For tap *n*, where *n* = 1 or 2, define

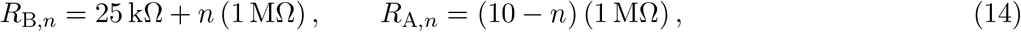

where *R*_B,*n*_ and *R*_A,*n*_ are the nominal resistances below and above the tap, respectively. For a multimeter with input resistance *R*_m_, the loaded division ratio is

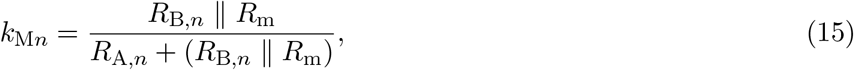

and the bank voltage is recovered from the meter reading as

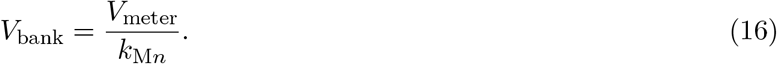

For a 10 MΩ multimeter input resistance,

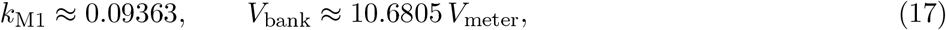

and

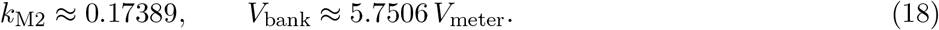

#### 5.2.8 Checkpoint A - Low-voltage discharge

This test verifies the discharge circuit’s topology at 24 V before any high-voltage work.

##### Setup

Connect a multimeter across the bank in voltage mode. Connect a 24 V bench PSU with a 0.5 A current limit directly across the bank, making sure to correctly connect the positive terminals together.

##### Static charge test

Turn on the PSU. Current should briefly rise, then fall to a few milliamperes as the bank becomes fully charged and the remaining current flows through the bleeder resistors. A continuous high current with the meter reading near zero indicates a polarity flip or a short somewhere; turn off the PSU and review the wiring. Once the bank holds voltage, disconnect the PSU from the charging circuit before turning it off. Leaving a turned-off PSU in circuit provides a secondary discharge path. Watch the multimeter for one minute. A drop of a few volts is expected from the bleeders. A fast decay indicates a short circuit.

##### Trigger test

Build the temporary trigger shown in Figure 7. With the bank charged to 24 V and the PSU disconnected, quickly press and release the push button. The bank voltage should snap to 0 V and a faint clicking noise coming from the TMS coil might be audible. If nothing happens, check the trigger polarity and pin assignment. For a different thyristor, you might have to adjust the resistance to deliver the required gate trigger current. Note that this trigger test is optional and provides a basic check that the thyristor works. A later test will cover this again with a proper gate driver (§5.3.5). The expected capacitor discharge trace is shown in Figure 7 as well.

**Figure 7:**
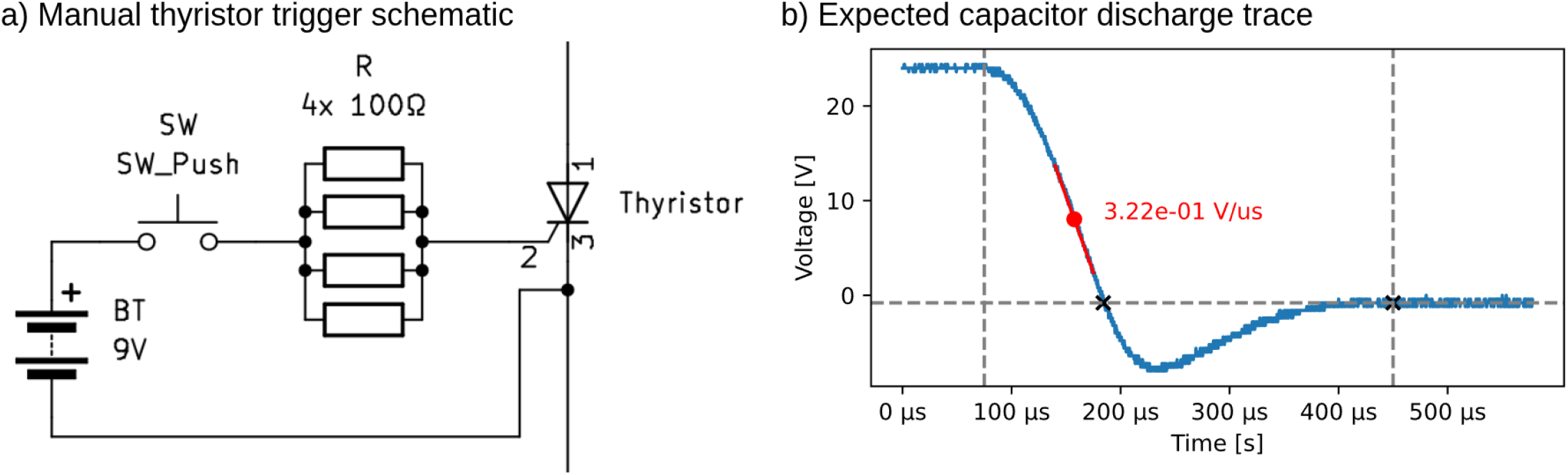
Checkpoint A test schematic and expected results. **a) Manual thyristor trigger schematic**. A temporary circuit to manually activate the gate of the thyristor. The push button should be quickly pressed and then released. This circuit is safe only for testing at 24 V capacitor charge. **b) Expected capacitor discharge trace**. Starting from a 24 V capacitor-bank charge, the voltage quickly drops and goes negative before all the energy is absorbed in the damping resistor. The positive lobe of the pulse should last between 80 and 100 µs with the previously mentioned capacitance and inductance. The 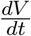 can be estimated by taking the slope in the middle of the positive lobe.

##### Estimate L

If you do not have access to an LCR meter, you can use this setup to estimate the inductance of your TMS coil. With oscilloscope probes placed across the capacitor bank terminals, extract the effective pulse length by placing a cursor at the beginning of the discharge pulse and a second cursor at the zero-voltage crossing. You should get a value of *t*_0_ between 80 and 100 µs. Rearranging Eq. (3), use

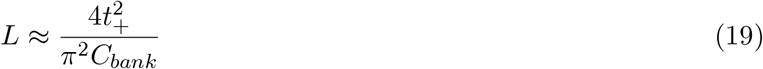

to estimate your coil’s *L*. For example, with our measurements of *t*_0_ = 90 µs and *C*_*bank*_ = 230 µF, we get *L* ≈ 14.3 µH which, compared to the measured 15.4 µH, is a 7.14% relative error. Note that this approximation assumes negligible parasitic resistance in the discharge circuit. Higher parasitic resistance and inductance will result in larger errors. The lossless equation theoretically overestimates the actual inductance of the system. Since the maximal operating voltage is proportional to the square root of the inductance (Eq. (9)), it will also be overestimated if using the approximate inductance found by this equation. If you estimate inductance using this method, keep in mind the large error it can introduce when computing your maximum 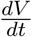 and other device limits. We recommend using a maximal voltage of no more than 80% of the calculated maximum voltage.

### 5.3 Control circuitry

An overview of the control circuitry is shown in Figure 8.

**Figure 8:**
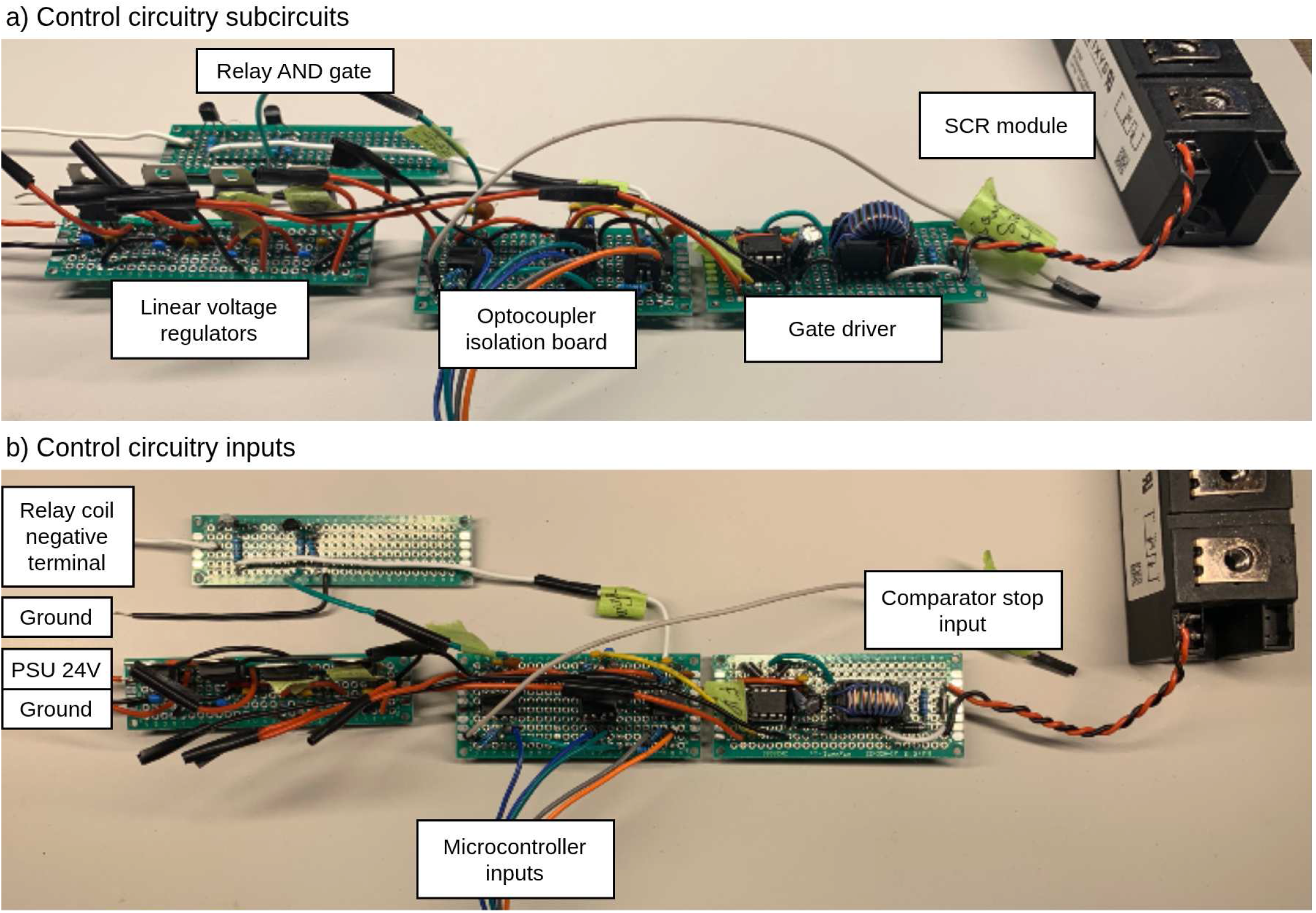
Overview of the control circuitry. **a) Control circuitry subcircuits**. All control-circuit perforated boards are shown with their respective labels. The SCR module is included in the photo since it receives a signal from the gate driver. **b) Control circuitry inputs**. All control-circuit perforated boards are shown with their respective inputs. The relay AND gate receives the negative terminal of the relay’s coil as input. The linear voltage regulators are connected to the PSU. The optocoupler isolation board receives different signals from the microcontroller and a signal from the measurement circuitry (`CMP STOP`) not yet introduced.

#### 5.3.1 Relay circuit

Build the relay AND gate on a perforated board. Wire the two NPN transistors so the relay coil energizes only when `CHARGE CTRL` is high and `CMP STOP` is low. Drive the relay coil from the 24 V PSU through the AND gate. Fit the antiparallel Schottky diode across the relay coil, and the 4700 µF electrolytic capacitor across the PSU terminals, taped to the relay. Leave each AND-gate input as a floating 22 AWG (0.64 mm diameter) lead for now; the optocoupler outputs in §5.3.3 will drive them. Figure 9 documents the relay setup and the AND gate wiring on the perforated board.

**Figure 9:**
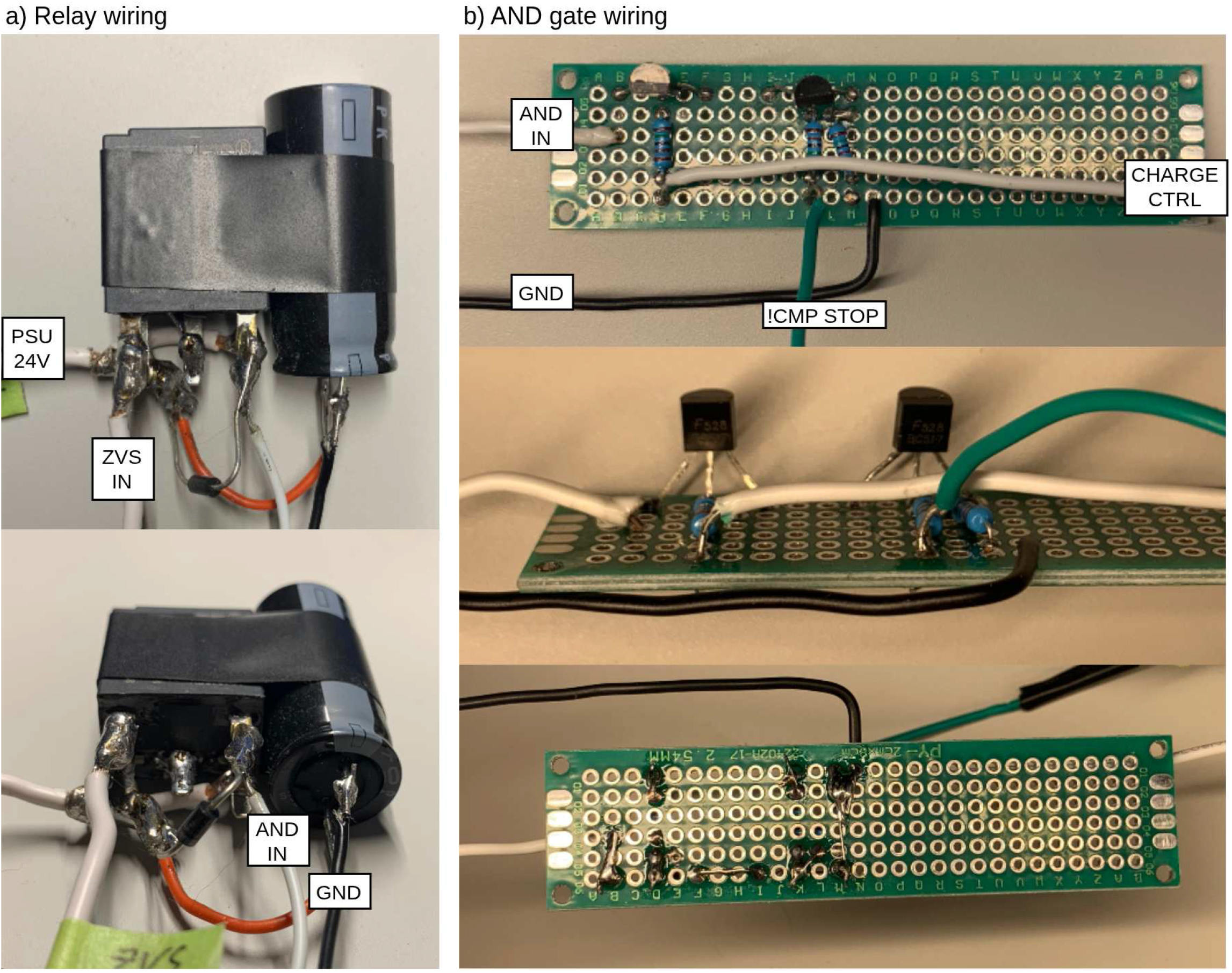
Wiring of the relay and the AND gate. **a) Relay wiring**. The relay is connected to the PSU and connects the PSU to ZVS IN when AND IN, the negative terminal of the coil, is grounded. A 4700 µF capacitor is connected across the terminals of the PSU and physically taped to the relay to help provide the initial power spike when the ZVS is activated. **b) AND gate wiring**. The AND gate connects AND IN to ground when `CHARGE CTRL` is high and `CMP STOP` is low. The `CMP STOP` signal comes from an optocoupler in an inverting configuration.

#### 5.3.2 Linear voltage regulators

On a separate perforated board, install one L7812CV (12 V) and one L7805CV (5 V) regulator, both fed by the 24 V PSU. Each regulator gets a 0.33 µF capacitor at its input to ground and a 0.1 µF capacitor at its output to ground. Use four capacitors in total. Do not share them between regulators. These rails are labeled PSU 12V and PSU 5V throughout.

#### 5.3.3 Optocoupler isolation board

On a third perforated board, install three Vishay VO2611 high-speed optocouplers carrying the `CHARGE CTRL, CMP STOP`, and `FIRE CTRL` signals from the microcontroller (an Arduino UNO R4 Minima in our build) to the power domain side. Use IC sockets to ease replacement after failures.

The three optocouplers fall into two configurations: `CHARGE CTRL` and `FIRE CTRL` are non-inverting; `CMP STOP` is inverting. The `CHARGE CTRL` output drives T1 in the relay AND gate, the inverted `CMP STOP` drives T2, and `FIRE CTRL` drives the gate-driver input in §5.3.4 (leave it floating for now).

Test the optocouplers by driving their inputs using the microcontroller with a fixed digital high or low and verify the output side with a multimeter.

#### 5.3.4 Gate driver

Build the gate driver around the Microchip TC4420CPA. This part is easy to destroy during debugging; buy several and use an IC socket.

The driver receives the `FIRE CTRL` signal from the optocoupler board and is powered from PSU 12V with two bypass capacitors (100 µF electrolytic and 0.1 µF ceramic) from VCC to ground to supply the initial current surge. The IC output drives the primary of a Pulse Electronics P0584NL 1:1 pulse transformer. Two Schottky diodes clamp the primary output node to PSU 12V and ground respectively, absorbing inductive kickback.

The secondary connects to the SCR gate via a series Schottky diode and a 10 Ω current-setting resistor; the secondary return connects the auxiliary cathode with an additional Schottky diode clamping the gate-to-cathode path. With approximately 12 V on the secondary, the resulting peak gate current well exceeds the SCR’s 0.5 A minimum. The instantaneous power in the 10 Ω resistor is around 15 W. Since manufacturers generally do not release instantaneous power ratings for 1/4 W resistors, it is difficult to know if a simple 10 Ω 1/4 W resistor is sufficient. In our tests, the resistor neither failed nor overheated since the pulses are extremely short (5 µs) and low frequency. For a larger safety margin, use four 40 Ω 1/4 W resistors in parallel.

Run the gate and auxiliary-cathode wires from physically adjacent points on the board and tightly twist them along the full run to the SCR pins. Parasitic inductance on this path distorts the pulse and can reduce the delivered 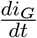.

The `FIRE CTRL` pulse must be 5 µs wide. The accepted tolerance interval is from 4.5 µs to 5.5 µs. This refers to the pulse measured across the gate resistor, not the microcontroller output, although both should be very similar. If the pulse is shorter, the SCR may not fully turn on and the gate region burns. If it is longer, the pulse transformer risks saturating (ET of 95 V µs) and the driver IC may fail. Verify the microcontroller pulse on a scope. Across multiple Arduino UNO boards, the `delayMicroseconds()` function was consistently long, and a programmed argument of 3 µs produced a measured 5 µs output. Calibrate the pulse length before connecting to the circuit.

##### Do not bypass

the optocoupler during debugging. A direct microcontroller connection to this inductive system risks an inductive kickback into the host USB port or computer.

Wiring for the linear voltage regulators, the optocoupler isolation board, and the gate driver is shown in Figure 10.

**Figure 10:**
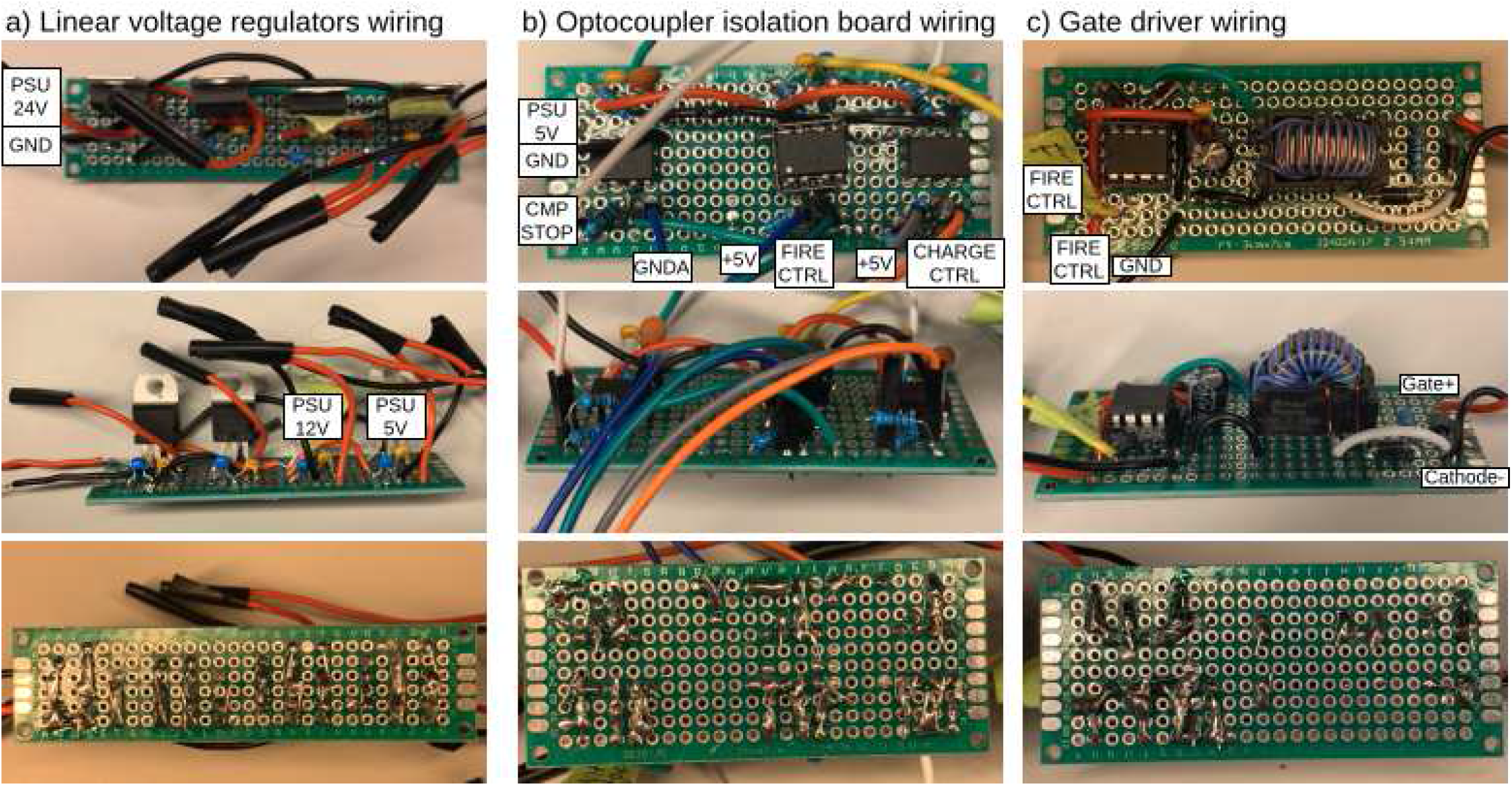
Wiring of control circuitry. **a) Linear voltage regulator wiring**. The board receives a 24 V input from the PSU and outputs two voltages. The board contains four linear voltage regulators, but only the right two are necessary for the TMS to function. The left two are unconnected. **b) Optocoupler isolation board wiring**. The board contains three optocouplers. Two non-inverting optocouplers transfer the `FIRE CTRL` and `CHARGE CTRL` from the microcontroller. An optocoupler in an inverting configuration transfers the `CMP STOP` signal received from the comparator circuit in the measurement circuitry. **c) Gate driver wiring**. The gate driver IC receives the `FIRE CTRL` signal from the corresponding optocoupler on the previous board. The signal is passed through a pulse transformer and sent to the gate of the thyristor.

#### 5.3.5 Checkpoint B - Gate driver test

This test estimates the SCR gate 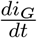 to ensure we are reaching the levels required by the module at high currents.

##### Setup

Confirm the microcontroller `FIRE CTRL` output is close to 5 µs on the scope. If your microcontroller does not produce 5 V logic high, adjust the optocoupler resistor and reference voltages.

##### IC test

Probe the gate-driver IC output during a single fire event. It should rise to approximately 12 V with a rising edge of a few nanoseconds.

##### Estimating gate 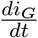

Using the same setup as Checkpoint A, but replacing the manual firing system with the gate driver, place the oscilloscope probe across the 10 Ω gate resistor. You may find it helpful to keep a second probe on the gate-driver IC output set as the trigger for the single-fire event. Use the curve to compute the rate of change of the gate current:

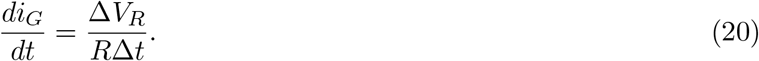

Measure Δ*V*_*R*_ and Δ*t* along the rising edge. We require 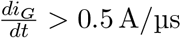.

Expected results are shown in Figure 11.

**Figure 11:**
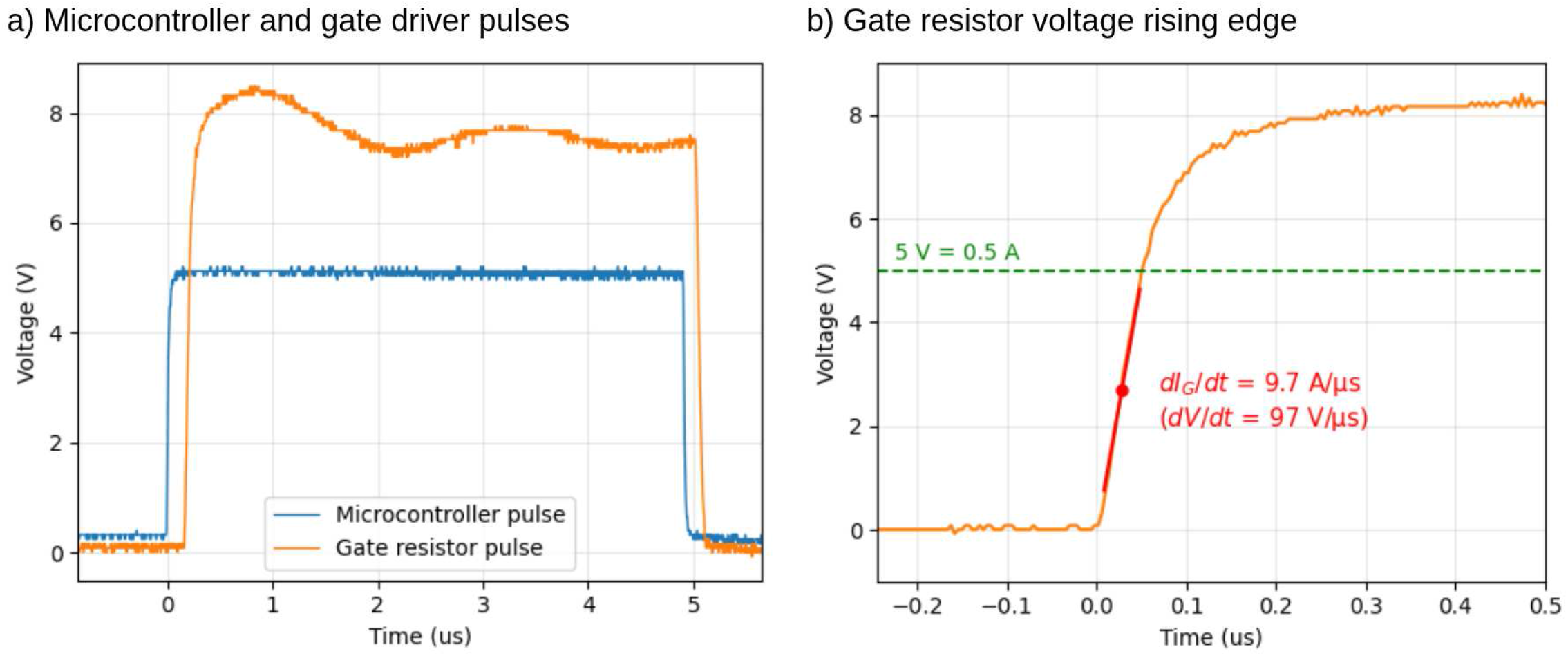
Checkpoint B expected test results. **a) Microcontroller and gate driver pulse**. The microcontroller pulse should be almost exactly 5 µs long to avoid saturating the pulse transformer and provide enough time for the thyristor gate to be properly activated. The pulse measured through the 10 Ω gate resistor should have a sharp rising edge. Since the gate driver IC is fed by a 12 V power supply, the voltage will rise above 5 V thereby providing more than the 0.5 A required by the thyristor gate. **b) Gate resistor voltage rising edge**. Zooming in on the voltage pulse across the 10 Ω gate resistor, we see that we reach 0.5 A in around 50 ns, resulting in an approximate 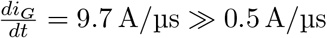.

Once this is achieved, you may proceed with the partial integration test described in §6.3.

### 5.4 Measurement circuits

#### 5.4.1 Voltage follower

Buffer the divider 1/401 (*V*_sense_) tap with a voltage follower stage before it feeds into the comparator and the linear optocoupler.

#### 5.4.2 Comparator circuit

Build the comparator on its own board around the LM393 dual voltage comparator IC. The trip threshold is set via the 5 kΩ potentiometer in a divider across the 9 V battery. Feed the buffered divider-tap voltage to one comparator input and the threshold to the other.

A second voltage follower buffers the comparator output before it drives a non-inverting optocoupler. The optocoupler’s control-domain output feeds both the microcontroller and a second optocoupler on the board described in §5.3.3; the latter gates the relay AND gate. The two-stage optocoupler path is necessary to preserve the three-domain isolation and protect the user and the equipment.

#### 5.4.3 Linear optocoupler

Build the linear optocoupler. Each photodiode is in a unity-gain op-amp stage with a matched 100 kΩ resistor. We use the UA741 on both sides. **Do not use** the two op-amps of a dual op-amp IC for the two sides. A dual op-amp package does not provide the galvanic isolation that the optocoupler does.

Wiring for the voltage follower, the comparator circuit, and the linear optocoupler are exposed in Figure 12.

**Figure 12:**
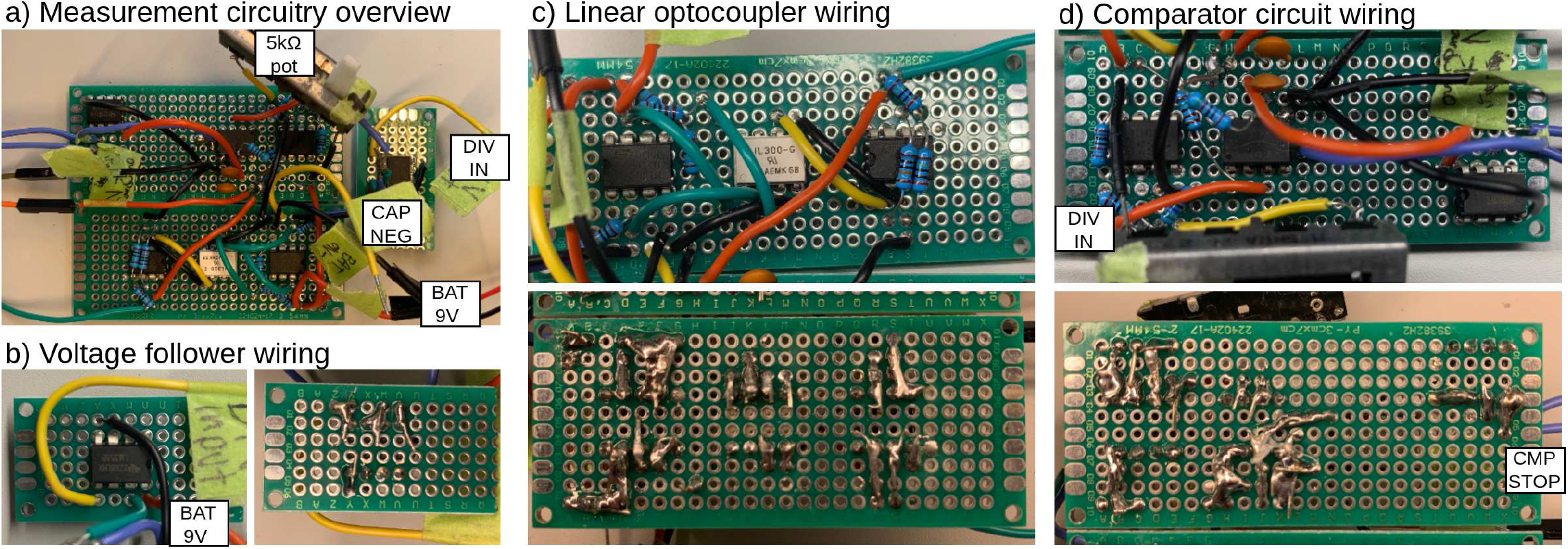
Wiring of measurement circuits. **a) Measurement circuitry overview**. All three perforated boards, containing the voltage follower, linear optocoupler, and comparator circuits. These circuits are all powered by a 9 V battery which shares ground with the capacitor bank. **b) Voltage follower wiring**. The voltage follower receives the voltage divider tap and provides a buffered version of the signal for the subsequent circuits to use. **c) Linear optocoupler wiring**. The linear optocoupler circuit transfers the analog voltage at the divider tap to the microcontroller. It uses the output of the voltage follower and produces the `CAP VOLTAGE` output. **d) Comparator circuit wiring**. The comparator circuit receives the divider voltage input from the voltage follower and sends a low or high signal based on a threshold set by the 5 kΩ potentiometer. The output of this circuit is fed to another voltage follower powered by the microcontroller-side voltage supply and the output of the voltage follower is sent to the microcontroller and to the `CMP STOP` optocoupler on the optocoupler isolation board.

#### 5.4.4 Checkpoint C - Measurement test

With the capacitor bank and PSU disconnected:

##### Comparator

Inject a DC signal below 5 V at the divider input. Sweep the potentiometer while probing the comparator output and the post-optocoupler output with a multimeter. The output should toggle when the threshold drops below the injected level.

##### Linear optocoupler

Inject a 0 V to 5 V 1 kHz sawtooth at the divider input and probe the post-optocoupler output with a scope, overlaying both traces. The output should track the input with negligible lag and no distortion at the upper end. The lower-voltage end may be slightly distorted because the op-amp negative rails are grounded. This is acceptable since the device does not need accurate low-end readings during operation. For accurate low-end readout, add a second 9 V battery for a −9 V negative rail on the high side, and a corresponding −5 V supply on the low side.

### 5.5 Charging circuit

Figure 13 provides an overview of the charging system.

**Figure 13:**
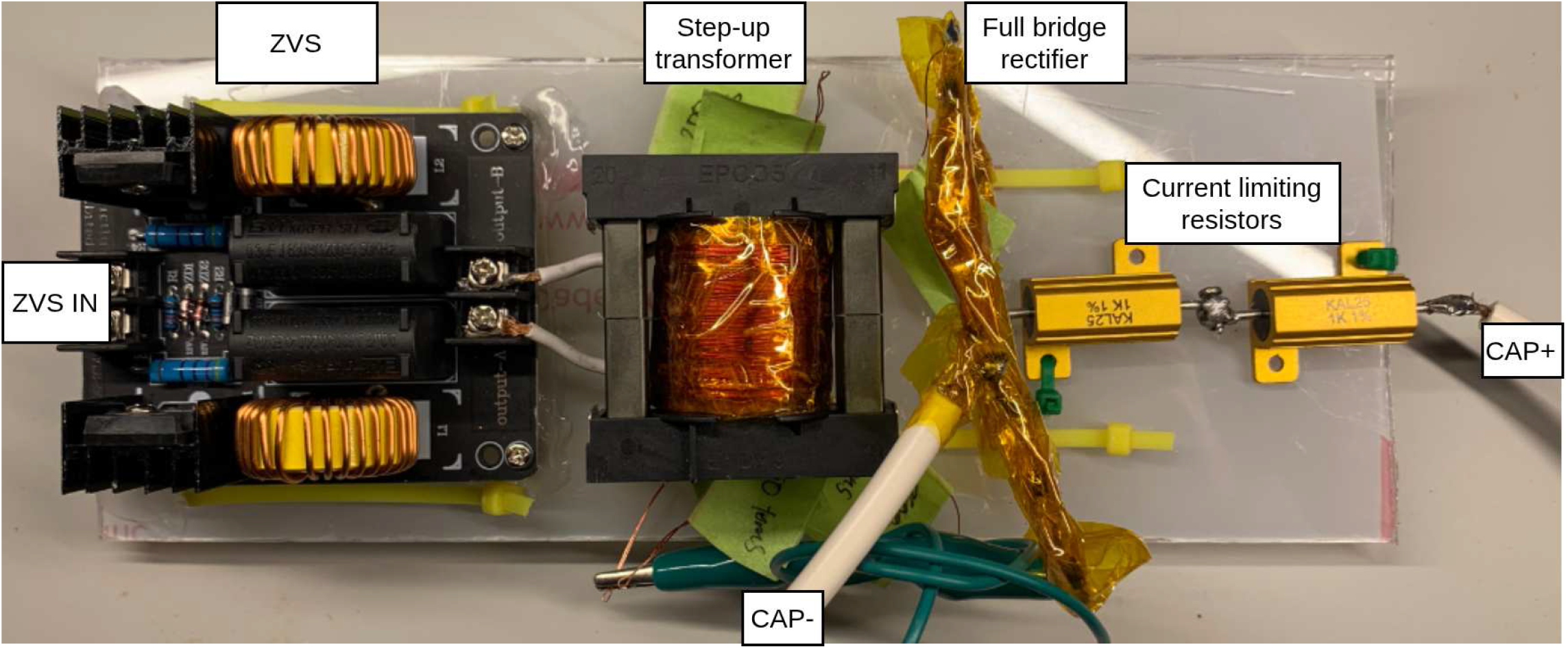
Overview of the charging circuit. The ZVS receives its input power through the relay. The output of the ZVS is fed into a step-up transformer with the secondary connected to a full-bridge rectifier. The output of the full bridge rectifier connects directly to the negative capacitor bank terminal and to the positive terminal via two 1 kΩ 25 W current-limiting resistors.

#### 5.5.1 ZVS driver

We use a commercial pre-built two-inductor ZVS driver. The selected driver has a 1000 W power rating, a maximum current of 20 A, and an acceptable input range of 12 V to 30 V DC. The two-inductor variant removes the need for a center tap on the transformer primary.

The ZVS requires an inductive load to oscillate. Defer testing until the transformer primary has been wound §5.5.2. With the primary connected, the ZVS oscillates the 24 V PSU input into a sine wave with peak voltage

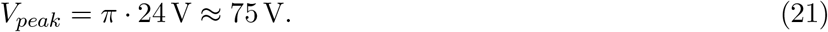

Treat the ZVS driver output as hazardous. Wear Class 00 or higher rubber gloves while handling the ZVS driver with an inductor at the output. When using the transformer, wear Class 1 or higher rubber gloves.

The ZVS driver has an initial power-demand spike that some bench PSUs might not be able to provide. Use the relay setup with the 4700 µF capacitor to turn on the ZVS driver while providing enough instantaneous power.

#### 5.5.2 Step-up transformer

The transformer uses the EPCOS/TDK B66368W1020T001 transformer bobbin, two ETD49/25/16-3C94 ferrite core halves, and B66368A2000X000 transformer clips.

##### Primary

Using 16 AWG (1.29 mm diameter) wire, wind 10 turns around the bobbin center. Leave sufficient lead length at each end to connect to the output terminals of the ZVS. Cover the windings with two layers of Kapton tape before winding the secondary. You may want to skip winding the secondary for now to test the ZVS driver. If doing so, make sure to insert the core halves as instructed in the paragraph labeled **Core**.

##### Secondary

Using 28 AWG (0.32 mm diameter) enameled copper wire, wind five layers of 50 turns each with a layer of Kapton tape between the winding layers. Take a tap at the end of each layer (50, 100, 150, 200, 250 turns), plus the initial 0-turn lead. Keep taps spatially separated to prevent arcing. Sand the protective coating from each tap with emery cloth before soldering.

##### Core

Insert one core half into the bobbin and flip the bobbin so that the ends of the inserted core are visible. To slightly reduce the inductance of the transformer, we create an air gap between the two core halves. Cut three small pieces of paper and place one piece on each leg of the inserted core half. We measured a gap of around 0.1 mm. Gently place the second half on top while making sure the pieces of paper do not move. Secure the core using the transformer clips. Figure 14 shows the process of winding the transformer.

**Figure 14:**
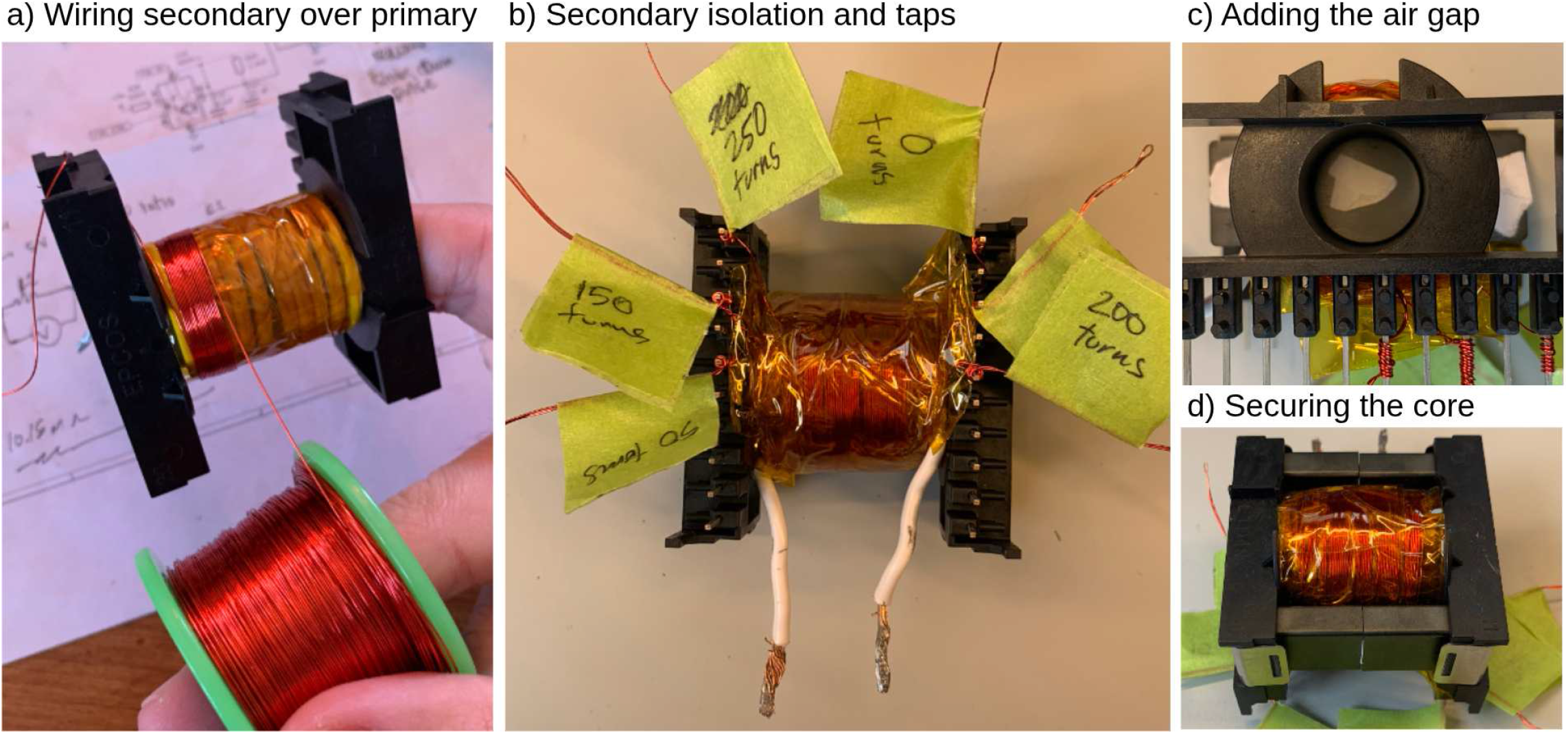
Winding the step-up transformer. **a) Winding secondary over primary**. The primary consists of 10 turns of 16 AWG (1.29 mm diameter) wire. Before winding the secondary, we place two layers of Kapton tape over the entire primary winding. The secondary consists of five layers of 50 turns each with 28 AWG (0.32 mm diameter) enameled copper wire. **b) Secondary isolation and taps**. After each layer of the secondary, we take a tap by extracting a small loop of wire outside of the transformer. The taps must not touch as there is a risk of arcing. Each layer of the secondary is covered in two layers of Kapton tape before winding the next one. **Adding the air gap**. To decrease the inductance of the transformer, we add small pieces of paper to each leg of a core half before inserting the second half. This enforces a small air gap in the core. **d) Securing the core**. The second half is finally placed in the bobbin and the transformer clips are installed to tightly hold the core in place.

At a primary peak of around 75 V, the tap peak voltages are approximately

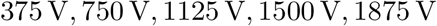

at the 50-, 100-, 150-, 200-, and 250-turn taps respectively.

#### 5.5.3 Full-bridge rectifier

The high-voltage rectifier uses four UF4007 diodes per leg, in series, for 16 diodes total; each rated at 1000 V repetitive reverse voltage.

Solder the four legs separately, then connect them into the standard full-bridge topology. Insulate every leg and joint with Kapton tape. The 0-turn lead of the step-up transformer’s secondary connects directly to the first AC terminal of the bridge. The second AC terminal is connected to a crocodile clip which is used to select the transformer tap before operation. The tap should never be changed during operation. If a change in tap is required, follow the full de-energization procedure. Wires with insulation that is not rated for high voltages should be viewed as bare metal wires. Whenever possible, add a few layers of Kapton tape to increase insulation.

The DC negative output connects via 18 AWG (1.02 mm diameter) HV silicone wire to the bank negative busbar. The DC positive output connects to the bank positive busbar via two 1 kΩ 25 W resistors in series (secondary-side current limiters) and 18 AWG (1.02 mm diameter) HV silicone wire. The rectifier output is electrically connected to the bank and is equally dangerous. Isolate all exposed metal with Kapton tape and ensure inaccessibility during operation.

Charging the capacitor bank to 1460 V implies 245 J are dissipated through the current-limiting resistors. At the tested interpulse interval of 30 s, this corresponds to time-averaged 4.1 W per resistor. This is below the unmounted power limit of the resistor. However, the initial instantaneous power is approximately 533 W per resistor. The chosen resistors must have a high overload rating on top of their continuous power rating. We choose not to mount the resistors since the periods for which they are used are short.

### 5.6 Safety hardware and enclosure

#### 5.6.1 Interlocks and charge-cutoff switch

Insert two interlocks and a charge-cutoff switch in series with the PSU output before the relay input:

- **A lid interlock** which is closed only when the enclosure lid is closed.
- **A dead-man interlock** which is held closed by a second person during testing.
- **A charge-cutoff switch** that disconnects the PSU feed.

The interlocks and charge-cutoff switch wiring is shown in Figure 15.

**Figure 15:**
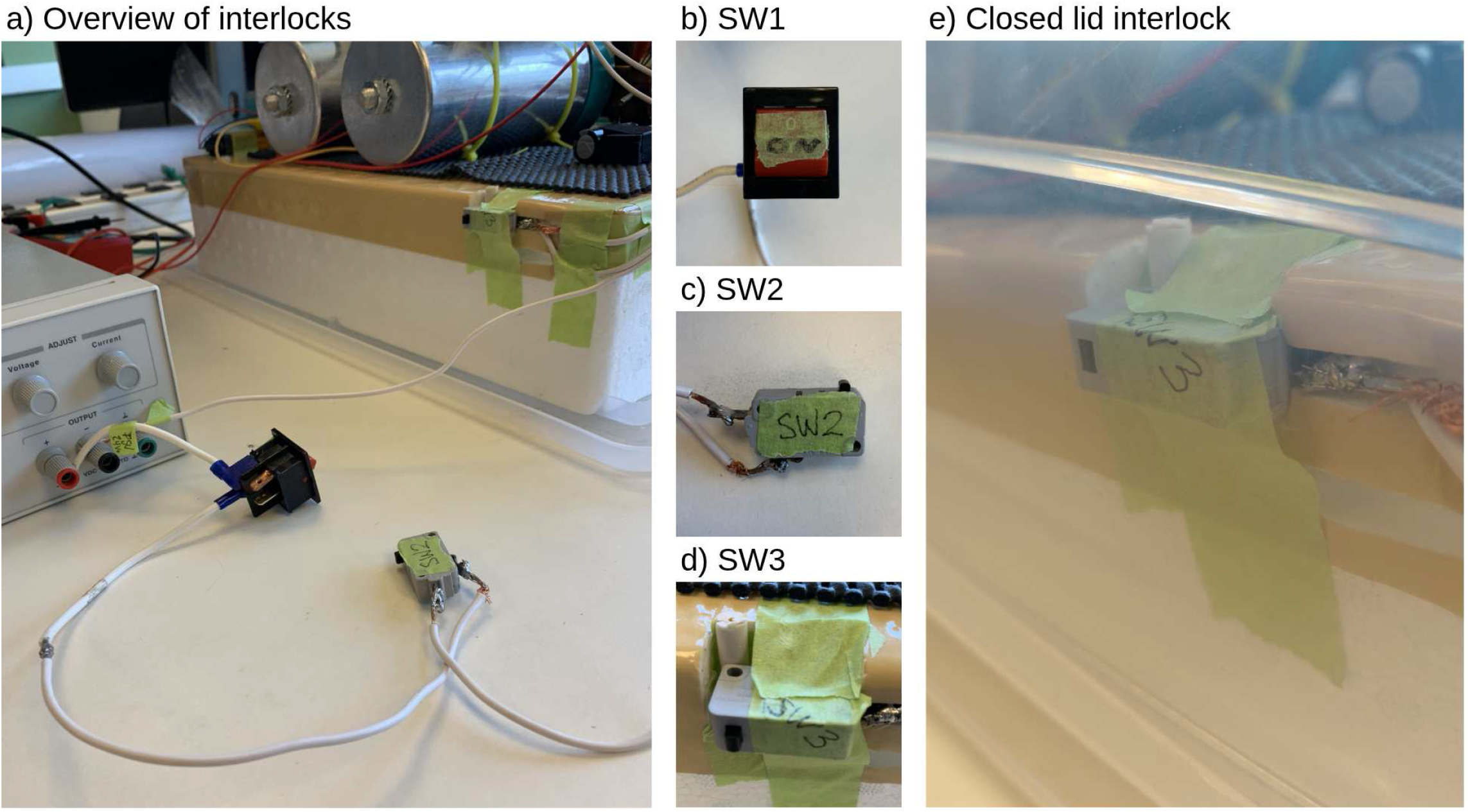
Interlock-wiring overview. **a) Overview of interlocks**. The PSU is connected to a large switch to turn off the power supply quickly in case of an emergency. The power also goes through two interlocks, one held closed by an operator and the other placed next to the enclosure so that it closes only when the lid is on. **b) SW1**. SW1 in the schematic is the charge-cutoff switch. **c) SW2**. SW2 in the schematic is the dead-man interlock switch. **d) SW3**. SW3 in the schematic is the lid interlock. **e) Closed lid interlock**. When the lid is positioned so that the high-voltage circuitry cannot be touched accidentally, the lid interlock closes, allowing the circuit to be energized.

These only halt charging and do not actively discharge the bank. After tripping, the bleeders slowly drain the bank. You may also use the safety discharge stick §5.6.3 for fast discharge.

#### 5.6.2 Enclosure and physical installation

We use a transparent plastic storage box as an enclosure. The transparent box allows the operator to spot sparks during testing. Small holes were made in the sides to allow control-domain wires to pass through. Similarly, for the TMS coil, a large hole was made at the front. The coil is connected to the discharge circuit and should therefore not be touched while the device is energized. The discharge circuit is mounted on a Styrofoam platform which isolates it from the box and the bench beneath. An adhesive mat on the Styrofoam keeps the capacitors and the SCR from shifting around. Zip-ties additionally secure components in place.

At maximal voltage, Lorentz forces are significant and can cause mechanical damage to your circuit. Pay particular attention to junctions where opposing currents separate, such as when the coil leads reach their respective busbars. Insert a non-magnetic separator at the Y-junction and apply Kapton tape under tension to oppose the forces.

A 12 V PC fan on the lid handles active cooling. The fan is powered by the 12 V linear voltage regulator and is connected using two 18 AWG conductors. The device was operated at the maximum capacitor-bank voltage of 1460 V (approximately 245 J), delivering 30 pulses over each 15 min test period at an interpulse interval of 30 s. Consecutive periods were separated by a 15 min cooling interval. No visible signs of thermal degradation were observed at this duty cycle, including during trials performed with the fan turned off. Because component temperatures were not instrumented, operation above this tested duty cycle is not recommended.

Figure 16 provides multiple views of the setup.

**Figure 16:**
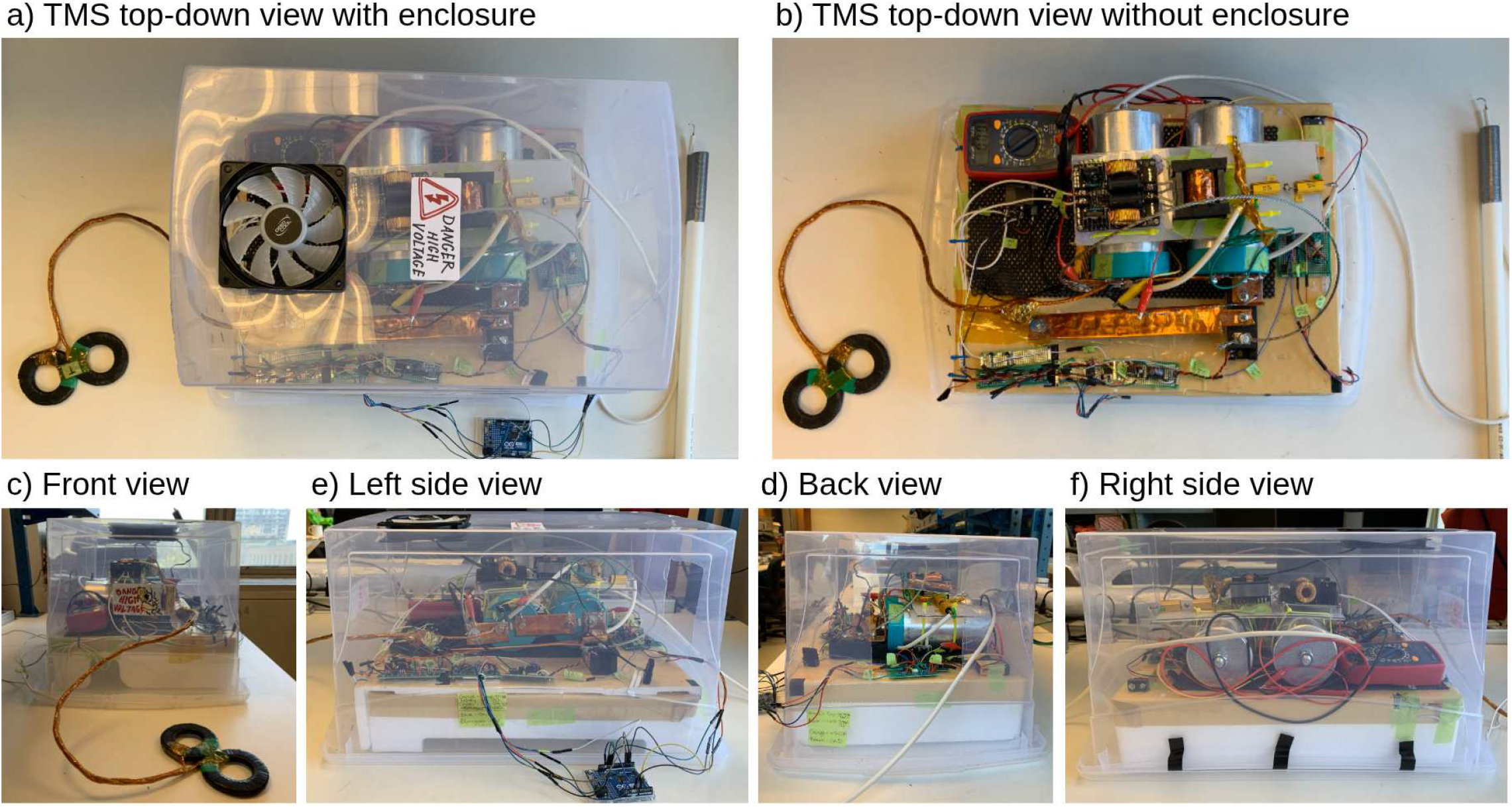
Enclosure overview. An overview of the enclosure used for the TMS. The plastic box allows the operator to see and detect any sparks or movement inside the box. Holes are made in the box to allow wires to pass through. A PC fan was installed on top of the enclosure to help with cooling.

#### 5.6.3 Safety discharge stick

A 1.5 m plastic pipe (PVC or PEX-B) carries an 18 AWG (1.02 mm diameter) HV silicone wire whose far end is shaped into a bare-wire hook and reinforced with solder as shown in Figure 17. The other end is connected to two 1 kΩ 50 W resistors in series, which are permanently wired to the bank negative busbar. A hole in the enclosure lets the silicone wire exit and the hook enter to touch the bank’s positive terminal without opening the box. Quick, firm contact minimizes sparking. A permanently connected multimeter indicates whether the bank has been properly discharged. In case the multimeter has accidentally been disconnected, the operator should connect the discharge stick for 10 seconds followed by leaving the device untouched for at least 20 minutes. This minimizes the chances of the capacitor bank still being charged. The operator can then reconnect the multimeter.

**Figure 17:**
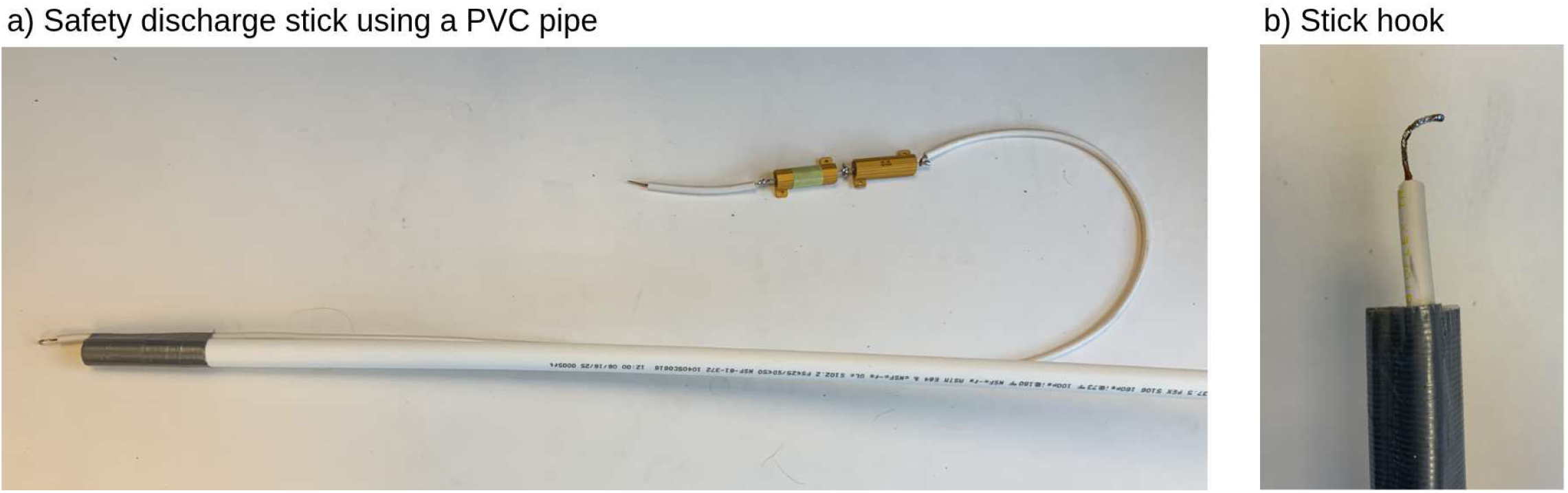
Safety discharge stick. **a) Safety discharge stick using a PVC pipe**. A PVC pipe is connected to the hook end of a high-voltage silicone wire. The other end of the wire connects to two energy-absorbing resistors which are themselves permanently attached to the negative capacitor bank terminal. **b) Stick hook**. The hook end connected to the stick is soldered to make it stiffer.

## 6 Operation instructions

### 6.1 Pre-operation safety

#### The voltages this device operates at can be lethal

The operator must wear Class 1 or higher insulating rubber gloves, impact-rated eye protection and safety shoes when the device is energized. Despite the protective personal equipment, direct contact with the system should never be made while it is energized. Place physical barriers between yourself and the high-voltage components, and keep a safe distance during charge, arm, and fire.

#### Never touch

the capacitor bank, the rectifier output, or any conductor electrically continuous with them while the circuit is energized or might still hold energy.

Place a multimeter across the bank for a live readout. The multimeter and its leads should be treated as an HV hazard at full bank voltage and should ideally be kept inside the enclosure. If your meter cannot read that high, probe across one 1 MΩ divider element for a ≈1:10 scaling (see §5.2.7) or use a 100x scope probe across the bank to measure the live voltage with your oscilloscope. Note that a regular oscilloscope probe shares the oscilloscope’s ground, bringing the high-voltage domain into the power domain.

A briefed second person must be present during operations. The enclosure lid must be closed, and the discharge stick connected.

Full safety checklists are available in the supplementary materials and in the project repository (Safety Checklists.pdf).

### 6.2 Software setup

Flash controller arduino.ino to the microcontroller. Install the Python TUI host app and its dependencies and then launch the app in a terminal window.

In the TUI, as shown in Figure 18, press P to list serial ports and select your microcontroller. The dashboard then shows live voltage readings and TMS status. If the status shows a hardware trip, press R to reset. If the trip persists, de-energize, raise the comparator threshold, and reset.

**Figure 18:**
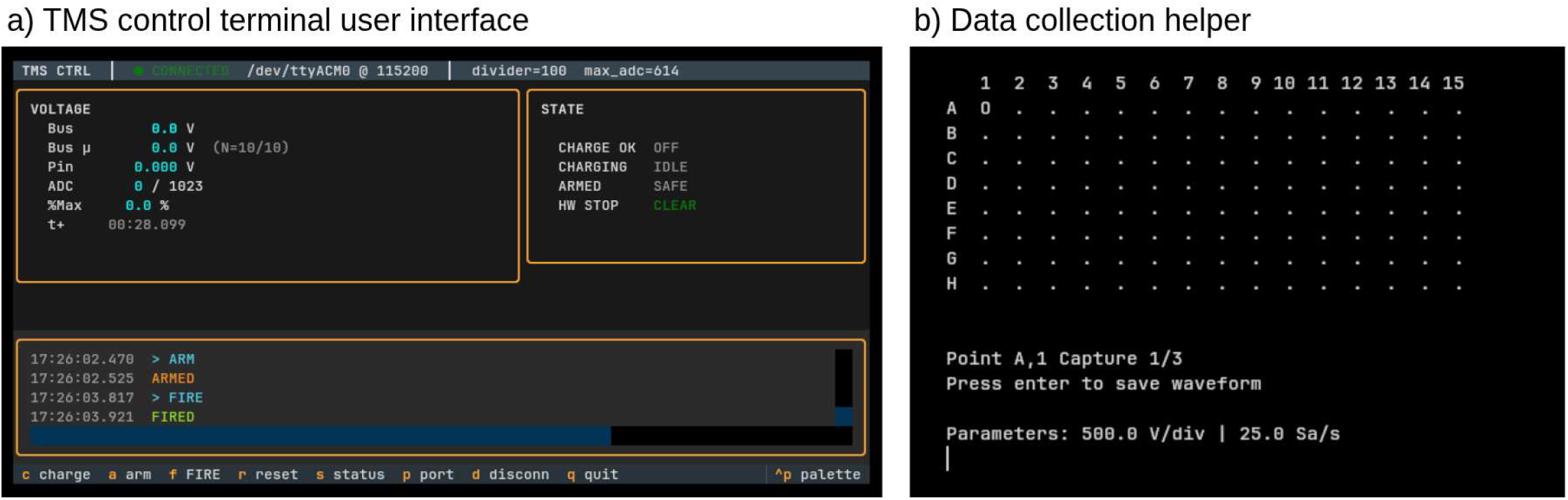
Host-side Python apps. **a) TMS control terminal user interface**. An interactive user interface that shows live changes in status to the user. The user can send commands via key presses which are then transmitted to the microcontroller via serial communication. The app also keeps a detailed log of past activity. **b) Data collection helper**. A second Python app communicates with the oscilloscope via USB using SCPI. It automates file naming and sets the oscilloscope to the proper data capture modes.

Table 4 provides more context on the different key presses and corresponding commands. Table 5 explains the possible asynchronous messages one might encounter looking at the serial port.

**Table 4:**
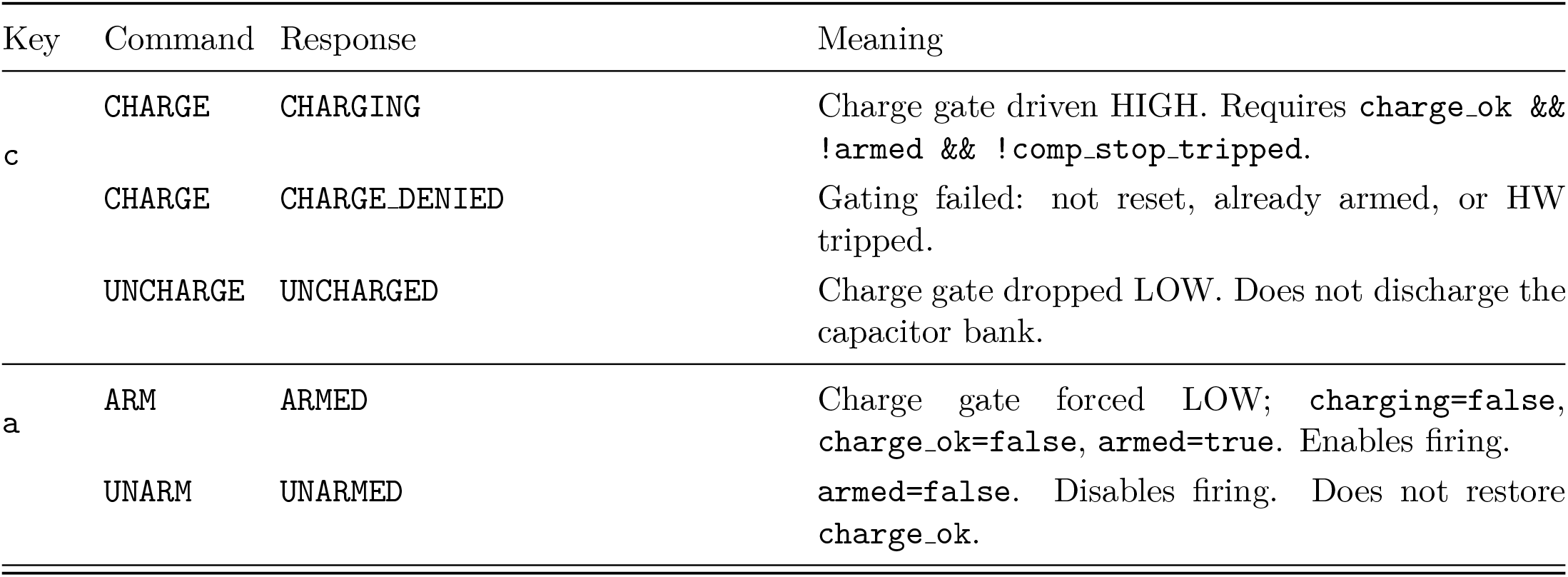
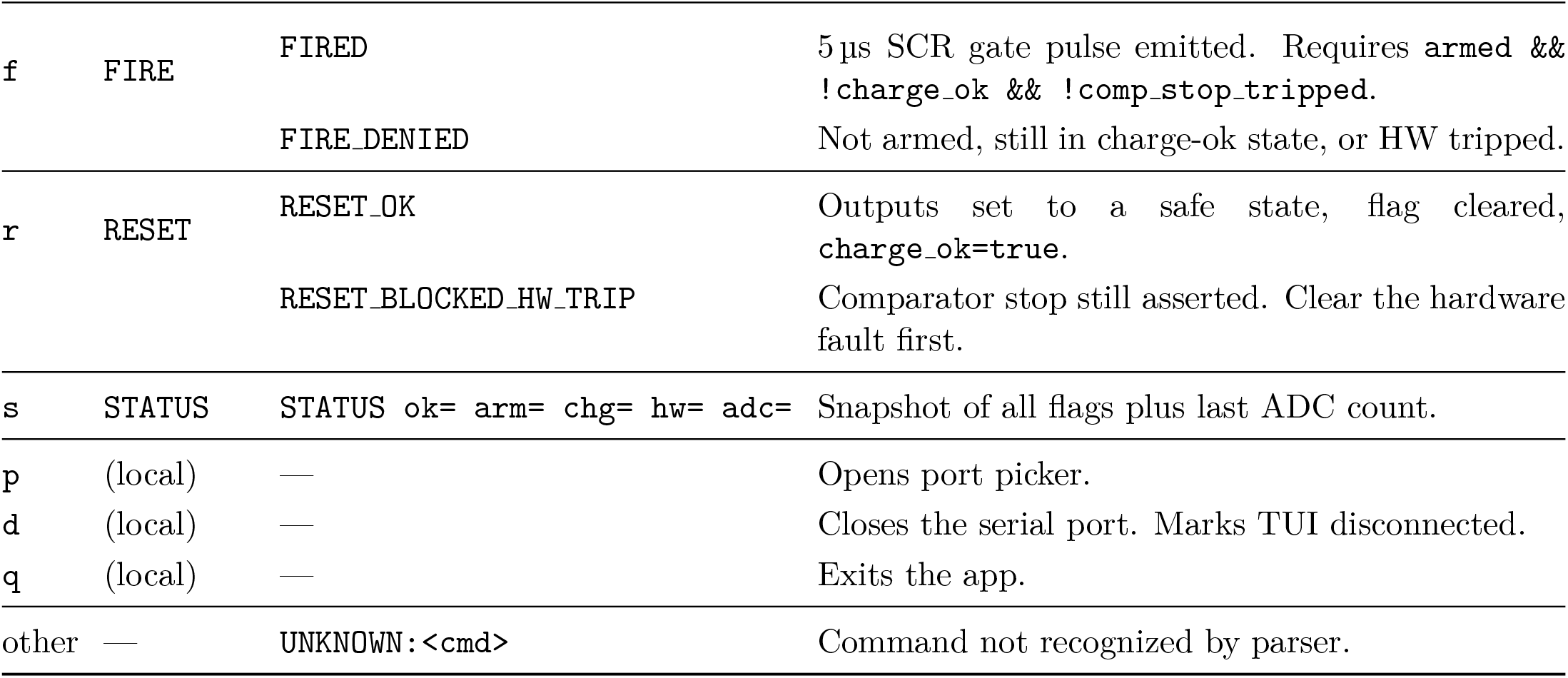
TUI keypresses, commands, and controller responses.

**Table 5:** Asynchronous controller messages.

| Message | Trigger | Meaning |
| --- | --- | --- |
| BOOT | Power-up / reset | Firmware started. |
| T t= adc= ... | 10 Hz | Telemetry line. Drives voltage and state boxes. |
| OVERVOLT | `adc_last > MAX_ADC` | Firmware voltage trip. |
| HW_STOP_TRIPPED | Comparator ISR latched | Hardware overvoltage comparator fired. Gate dropped in ISR, flags cleared in loop. |

### 6.3 Low-voltage test

Disconnect the relay output from the ZVS input and wire it directly across the terminals of the capacitor bank as shown in Figure 19. This bypasses the charging circuit and lets the relay charge the bank to 24 V directly. Use this run to practice the safety checklist and remote-controlled functions of the device.

**Figure 19:**
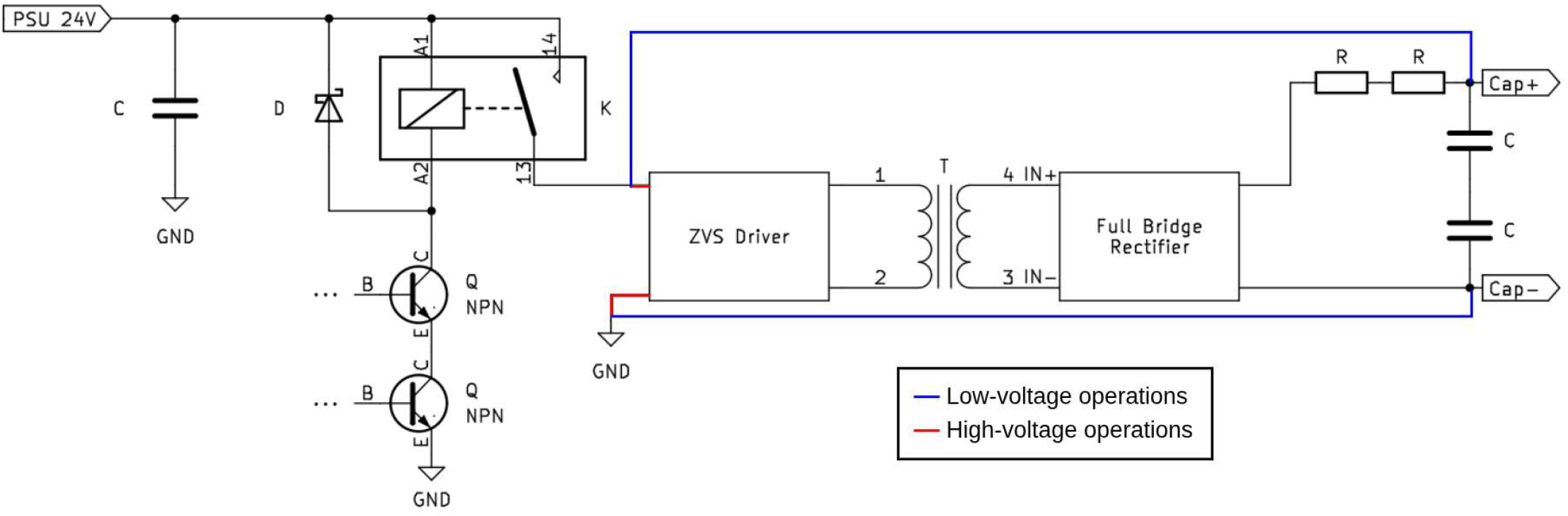
Charging circuit diagram for low- and high-voltage tests. For low-voltage tests, the output of the relay is sent directly to the capacitor bank terminals. Represented in the diagram by the blue wires. For the high-voltage tests, the output of the relay goes to the ZVS driver which is connected to the capacitor bank via the high-voltage charging chain. Represented in the diagram by the red wires.

1. Turn on the PSU. Quiescent current draw should be under 1 A.
2. Press `r` to reset; verify the dashboard reads nominal.
3. Press `c` to charge; the bank should reach 24 V quickly.
4. Press `a` to arm.
5. Press `f` to fire. Capacitor voltage should snap to 0 V. A faint coil click might be audible.
6. Turn the PSU off.

To test the comparator and linear optocoupler circuits at this voltage, switch the divider to the *V*_2M_ tap and adjust the potentiometer accordingly. The divider ratio should also be adjusted in the TUI app at launch for the voltage readings to match actual voltages. Use the `--divider` flag to set your divider ratio.

### 6.4 High-voltage test

Reconnect the relay output to the ZVS input. Move the full-bridge rectifier AC input to the 50-turn tap (with the 0-turn lead on the other bridge AC terminal). Always fully de-energize before changing a tap. This yields a bank charge voltage of approximately 375 V, which can be lethal. All high-voltage safety guidelines apply.

1. Confirm pre-charge safety state using the checklist.
2. Press `r` to reset.
3. Press `c` to charge. The ZVS may draw a startup spike of around 10 A. Set the PSU current limit accordingly. A faint ringing noise coming from the ZVS may be audible. Current draw falls as the bank charges.
4. Press `c` again to stop charging.
5. Monitor the multimeter for a minute. The voltage will drop by a few volts every few seconds due to the bleeders. Faster decay might indicate a short circuit. If that is the case, abort and follow the safe de-energization procedure using the de-energize checklist before debugging.
6. Press `a` to arm. This stops charging.
7. Press `f` to fire. Voltage should snap to 0 V. A distinct TMS coil click should be audible at this voltage.
8. Safely turn the device off following the safety checklist.

If 375 V succeeds, repeat the previous steps at higher taps. As voltage rises, Lorentz forces on internal conductors become significant. Watch for visible motion or sparking through the transparent enclosure.

#### Maximum voltage

With the bank and coil specified here, **do not exceed** a capacitor voltage that would exceed the bank’s 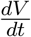 rating of 24 V/µs. Revise this limit only if you change the coil inductance or bank capacitance and have re-derived the discharge 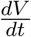.

If the bank ever exceeds the target voltage, either wait for the bleeders to slowly discharge the capacitors or proceed with manual discharge following the proper safety checklist.

We recommend capturing the bank discharge curve on the scope at each new voltage using a 100x probe across the bank, and deriving the peak 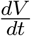 to confirm it remains within the bank rating as shown in Figure 20.

**Figure 20:**
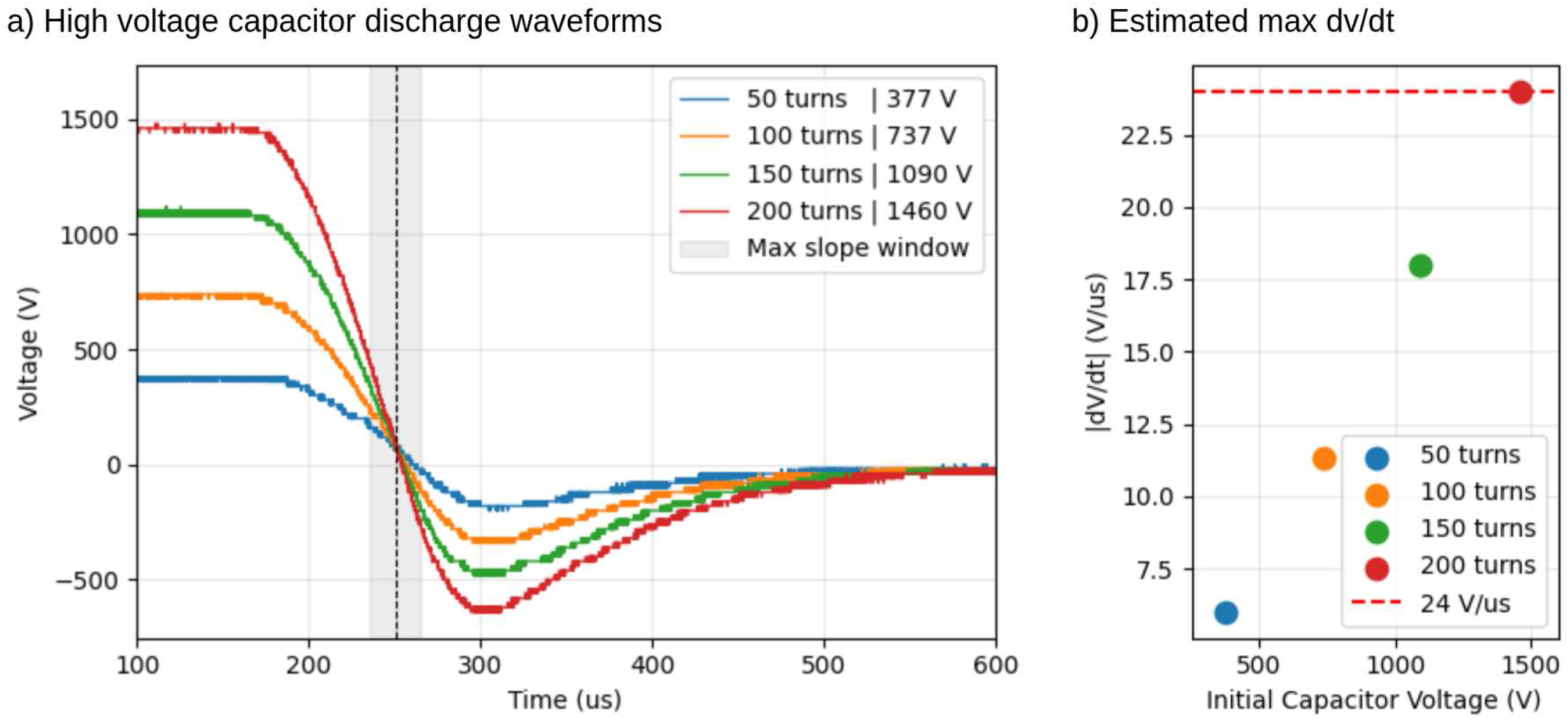
Capacitor discharge waveforms and dv/dt calculation. **a) High voltage capacitor discharge waveforms**. The plot shows the capacitor-bank discharge waveform for the 50-, 100-, 150-, 200-turn taps. A 30 µs window is taken around the area of maximal change in voltage to derive the maximal 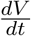. **b) Estimated max dv/dt**. Maximal 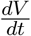 is derived from the voltage drop across the 30 µs window for all curves in a). The highest 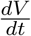 does not exceed 24 V/µs.

## 7 Validation and characterization

We characterize the device by reconstructing the peak coil 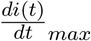 from external 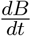 measurements, then feeding the recovered 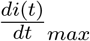 into a SimNIBS head model to estimate the induced cortical *E*-field at two standard stimulation sites. This idea of measuring the 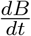 on a surface near the coil has previously been used and validated by a few groups [18, 19]. Drakaki et al. [19] validated their measurement setup by comparing their estimated 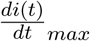 with the value reported by the stimulator. They report such data points for the MagVenture MagPro X100 with the MagVenture MRI B91 coil. We test our measurement methodology on the same stimulator-coil combination in §7.7 to validate that our error is within a range similar to that found by Drakaki et al. They saw a stimulator-reported 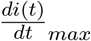 of 203 A/µs while they calculated 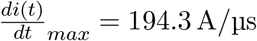.This corresponds to a 4.3% error normalized by the stimulator-reported 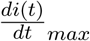.

### 7.1 Pickup coil

Wind 10 turns of 28 AWG (0.32 mm diameter) enameled wire tightly on a 3D-printed cylindrical former of 1 cm outer diameter ([pickup_cylinder.FCStd](https://data.mendeley.com/datasets/gb8tgtfk3c/2/files/8c01c343-8385-4d5d-b375-c3e3a92392a1)). The leads exit the coil together and are twisted tightly together to suppress stray inductance. Fix the windings in place with a few drops of glue and cut the leads to a 15 cm length.

After the glue cures, trim away as much of the PLA former as possible to help with later manipulations. PLA is non-magnetic, and perturbations of the field due to the leftover PLA are expected to be negligible. Trimming is done mainly to improve ease of use.

### 7.2 Measurement and positioning jig

Print the supplied PDF file which contains an 8 × 15 cm grid ([TMS_field_grid.pdf](https://data.mendeley.com/datasets/gb8tgtfk3c/2/files/9c4c7682-9170-453c-8176-e762fff8f7a5)). Disable your printer’s auto-scaling setting before printing the grid and verify the grid dimensions are accurate once printed.

A 3D-printable pickup coil positioning guide is available on the project repository as well ([sensing_cross.FCStd](https://data.mendeley.com/datasets/gb8tgtfk3c/2/files/b43b646b-9718-44cc-8c03-0421bcf9fffd)). The guide lets you slide the coil beneath the TMS coil and rotate it on all three axes to sample each axis at the same coil-to-coil distance.

Place the TMS coil on a sheet of acrylic glass over the printed grid, elevated on uniform-height supports (dice, printed legs) so the pickup coil can pass underneath. Record the coil-to-coil distance for later analysis.

Images of the pickup coil and of the positioning jig are provided in Figure 21.

**Figure 21:**
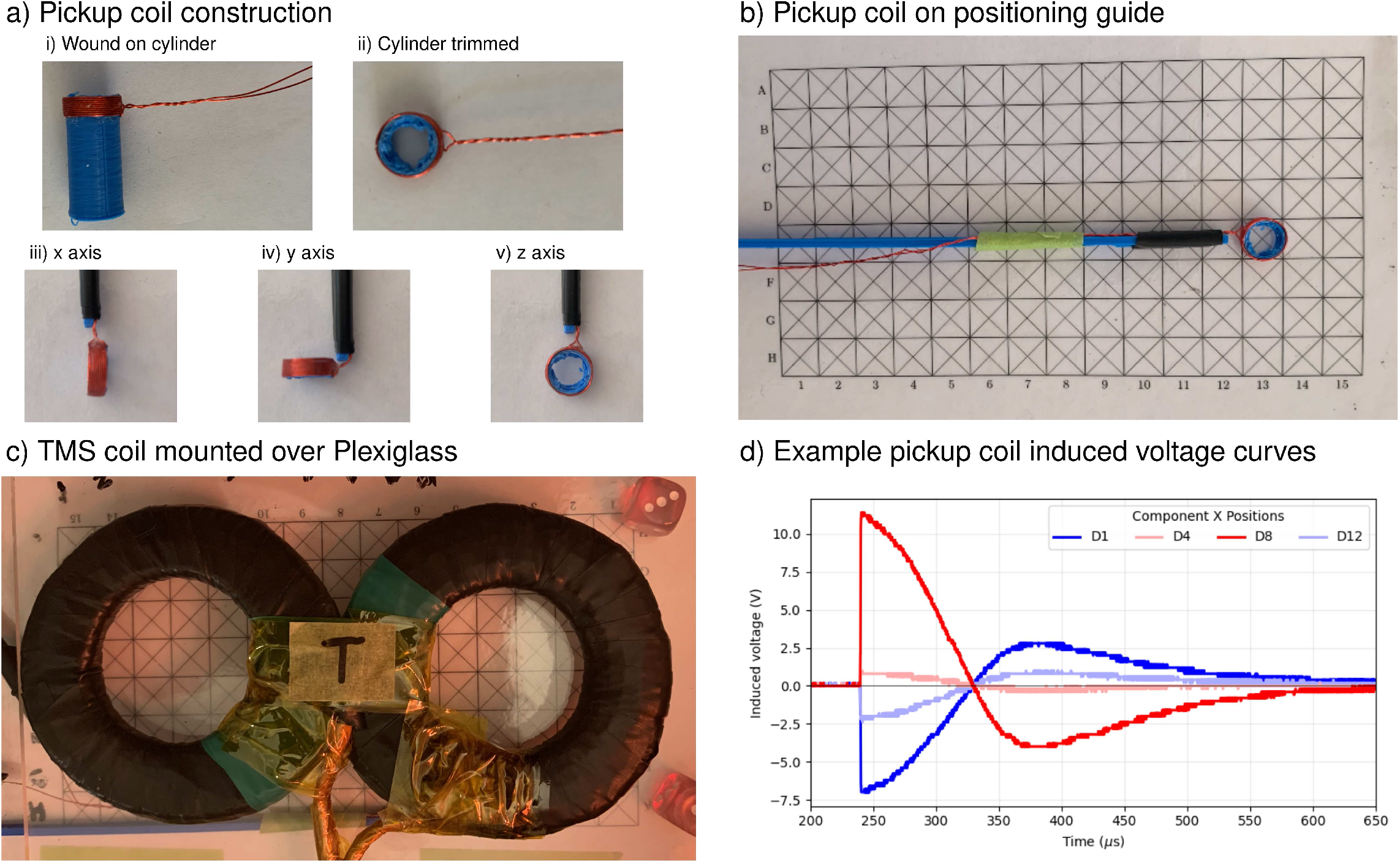
Pickup coil and measurement setup. **a) Pickup coil construction**. 28 AWG (0.32 mm diameter) enameled wire is wound 10 times around a 3D-printed cylinder and fixed in place with a few drops of glue (i). The plastic cylinder is then trimmed (ii). The pickup coil can be adjusted to capture different components of the field (iii-v). **b) Pickup coil on positioning guide**. A 3D-printed positioning guide allows the user to manipulate the pickup coil at some distance to more easily move it when under the TMS coil. Behind the guide and the pickup coil is an 8 × 15 cm printable grid which helps position the pickup coil during data collection. The grid has the same dimensions as the TMS coil. **c) TMS coil mounted over Plexiglass**. The TMS coil is slightly elevated from the positioning grid to allow the pickup coil to slide under. Using a Plexiglass sheet makes it easier to place the pickup coil at the correct position. **d) Example pickup coil induced voltage curves**. Four example pickup coil induced voltage curves from the X component at capacitor bank voltages of 1460 V. The displayed curves come from the positions D1, D4, D8, and D12. The maximal value reached by the induced voltage curve is used for deriving the 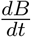.

### 7.3 Oscilloscope settings for 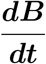 measurements

Connect the pickup coil leads to CH1 via a probe and the 100x bank-voltage probe to CH2. Note that this connects the high-voltage domain to the power domain. Use:

- CH1: 50 mV/div at 24 V — 5 V/div at 1500 V
- CH2: 5 V/div at 24 V — 500 V/div at 1500 V
- Time base: 25 µs/div
- Trigger: CH2, falling edge, at 1/2 the charge voltage. Use single-shot capture mode.

Upon firing, you should see two traces: the capacitor-discharge curve, as shown in Figure 7 b) and in Figure 20, and the pickup-voltage spike, as shown in Figure 21 d).

### 7.4 Data acquisition procedure

We use the Siglent SDS 1052DL+ oscilloscope connected to a computer via USB Type-B. A Python data-acquisition script (collect grid.py) automates capture, file naming, and per-shot scope reset. The script targets Siglent’s SCPI command set and should adapt to other Siglent models without modification and to other vendors with minor command-string changes.

At 24 V bank charge, we sample three pulses per grid point on each of the three axes, mapping the full topography of each 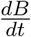 beneath the TMS coil. At higher bank voltages, we sample 30 strategically chosen grid points on the x and z axes and 32 on the y axis. Those grid points are marked in Figure 22. The points were selected from regions of greatest topographic variation to include both large- and small-magnitude measurements.

**Figure 22:**
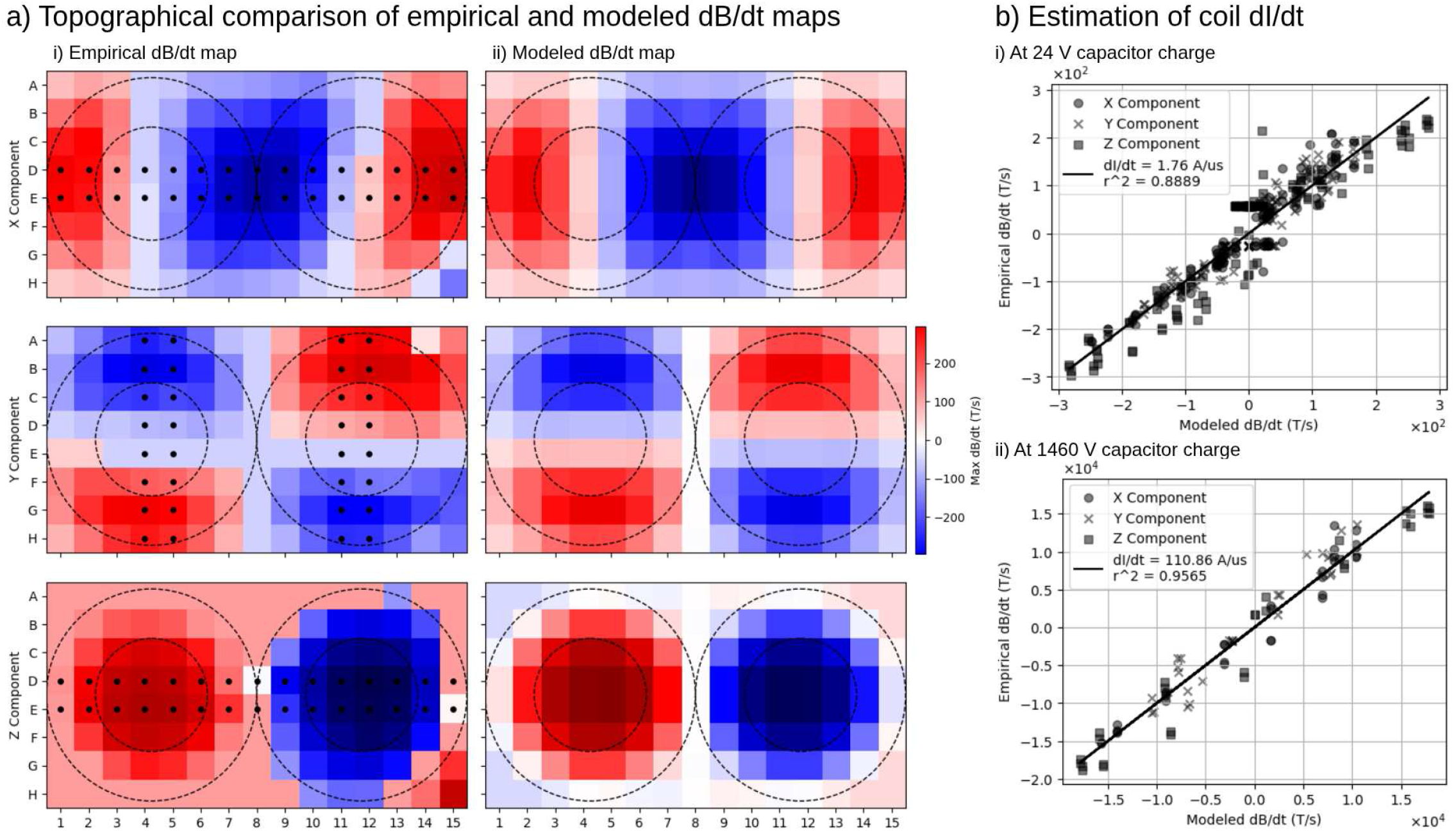
Estimating coil dI/dt from dB/dt topographical maps. **a) Topographical comparison of empirical and modeled dB/dt maps**. The empirically derived dB/dt topographical map obtained from probing the field generated by 24 V pulses using the pickup coil (i) closely matches the modeled dB/dt map (ii), which uses a simplified coil model. The grid points marked correspond to the grid points sampled during high-voltage testing. **b) Estimation of coil dI/dt**. A least-squares regression is performed to find which dI/dt produces the topographical map most similar to the empirical one. With a 24 V capacitor-bank voltage (i), we find a value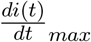 of 1.76 A/µs gives us a *r*^2^ = 0.8889. With a 1460 V capacitor-bank voltage (ii), we find a value 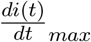 of 110.86 A/µs gives us an *r*^2^ = 0.9565.

### 7.5 Estimating 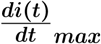

1. A fourth-order digital Butterworth low-pass filter with a cutoff frequency of 120 kHz was applied to the data. The filter was applied in the forward and reverse directions using `scipy.signal.sosfiltfilt`, producing zero phase shift and an effective eighth-order magnitude response. For each spatial point at each voltage, we extracted the absolute maximum and average over trials. Since we know our pickup coil has *NA* = *N* ·*πr*^2^ = 10·*π*(0.5×10^−2^)^2^, we recovered the field rate of change via Faraday’s law:

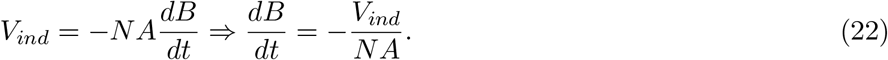

We model the TMS coil as two layers of five concentric circular loops per wing, mirrored to form the figure-of-eight geometry. This corresponds to twenty circular loops. For each measurement location *i*(*t*), the Biot-Savart model returned the signed magnetic-field component along the pickup-coil normal per unit coil current. Because the magnetic field is linear in coil current, the same coefficient relates the field derivative to the current derivative. The measured peak field derivative *y*_*i*_ is modeled as *y*_*i*_ = |*g*_*i*_|*k* + *ϵ*_*i*_, where *k* is the peak coil current derivative in amperes per second and *ϵ*_*i*_ is the measurement and modeling residual. *g*_*i*_ may be interpreted as (*dB*_*i*_*/dt*)/(*dI/dt*).

Measurements from the x-, y-, and z-oriented pickup coils were pooled into one regression. We used an unweighted least-squares fit constrained through the origin, so that

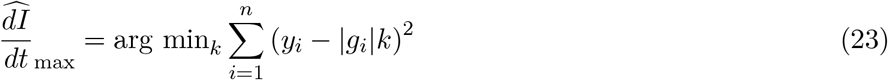

Figure 22 provides the 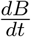 empirical and theoretical maps and context on the linear regression.

At 24 V, we found a 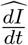 with *r*^2^ = 0.8889. At 1460 V, we found a 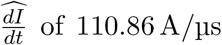 with *r*^2^ = 0.9565.

### 7.6 Cortical E-field estimation

We feed the recovered 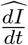 into SimNIBS 4.6.0 [12] on the Ernie example head model. Using the SimNIBS Python API, we wrote a custom script (create_fo8_coil.py) to recreate our self-wound coil in SimNIBS. This allows us to better approximate the cortical *E*-fields. We modeled the coil with a 0.706 mm physical standoff to account for the thickness of the Kapton and electrical tape used to insulate the coil. In SimNIBS, the coil was placed directly on the scalp with a skin distance of 0 mm and the default coil orientation was kept.

At Oz (occipital, visual cortex) and C3 (motor cortex), with 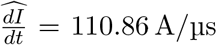, SimNIBS reports (Figure 23):

**Figure 23:**
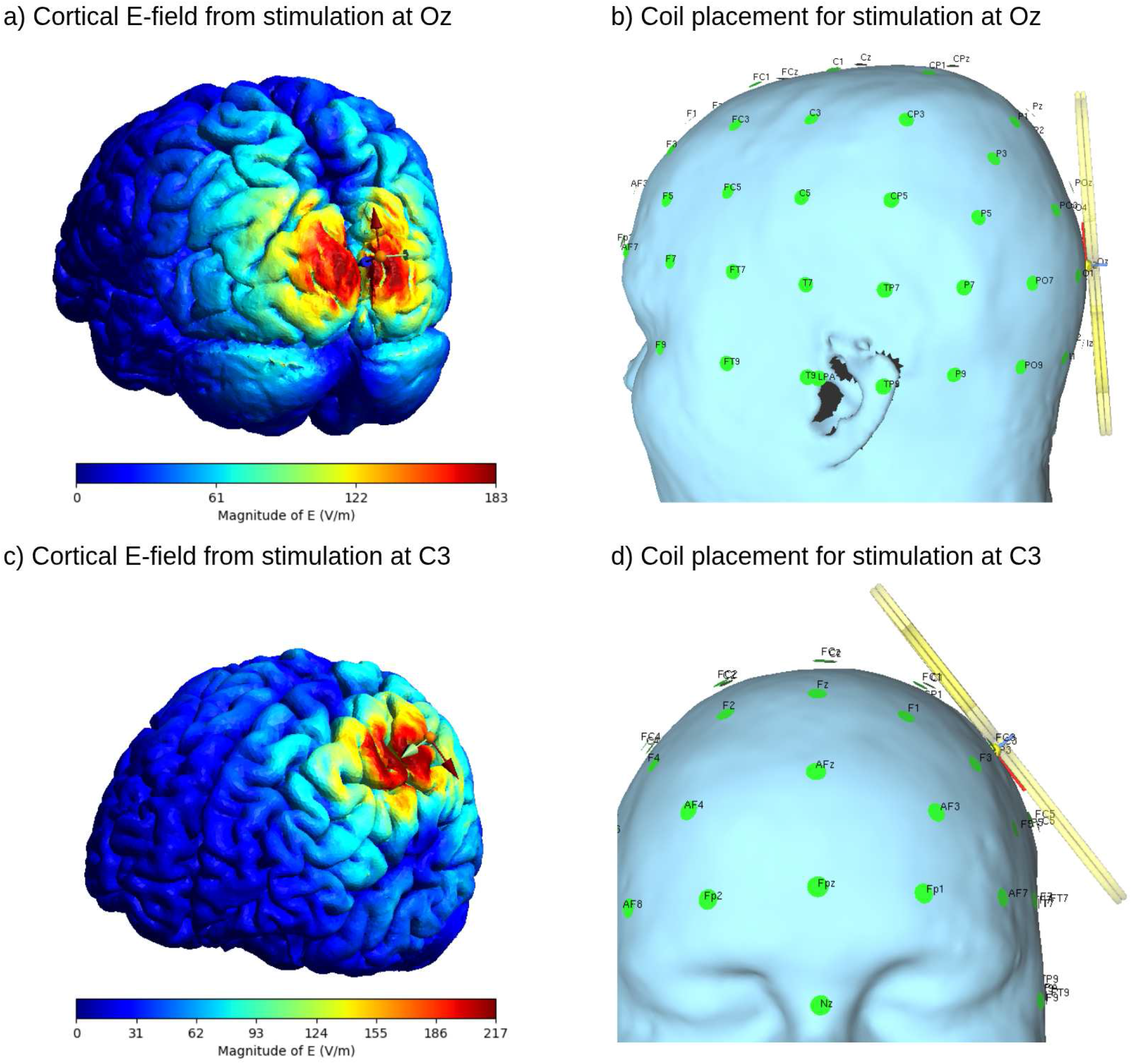
Cortical *E*-fields from SimNIBS simulation. **a) Cortical** *E***-field from stimulation at Oz**. The *E*-field magnitude plotted on an example anatomy. Our custom self-wound coil was placed at the Oz reference on the Ernie example anatomy in SimNIBS. A maximum *E*-field magnitude of 183 V/m, with the 99.9th percentile at 159 V/m, was estimated with a coil 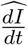. of 110.86 A/µs. **b) Coil placement for stimulation at Oz**. On the example anatomy, the coil, seen in yellow, is placed directly on the scalp surface at the Oz landmark. **c) Cortical** *E***-field from stimulation at C3**. The same simulation but this time with the coil placed at the C3 reference point gave a maximum *E*-field magnitude of 217 V/m, with the 99.9th percentile at 196 V/m. **d) Coil placement for stimulation at C3**. On the example anatomy, the coil, seen in yellow, is placed directly on the scalp surface at the C3 landmark.

- 99.9th percentile of cortical *E*-field at Oz: 159 V/m.
- 99.9th percentile of cortical *E*-field at C3: 196 V/m.

For context, Danner et al. [20] reported a mean modeled motor threshold, expressed as a cortical electric field, of 120 ± 24 V/m using a monophasic pulse waveform. Because no study provides a value directly comparable to our output, we can only provide the following comparison as general context rather than as a complete comparative analysis.

Before correcting for any differences in methodologies, this figure is approximately 63% of our stimulator’s maximal output. Although they do not state the effective pulse length of their monophasic stimulation, we find in the eXimia TMS system’s documentation that it produces a 70 µs effective pulse length when paired with the Focal Monopulse 8-Coil.

To correct for the difference in pulse length between their stimulator and ours, we use Lapicque’s relation in Eq. (1) with *τ*_*m*_ = 196 µs [16] and find the ratio

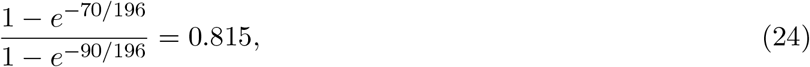

meaning their 120 V/m figure corresponds to 97.8 V/m when using our stimulator. At maximum output with the coil at C3, our stimulator produces an estimated cortical *E*-field of 196 V/m, twice the Danner et al. motor-threshold estimate.

Finally, we cannot present these numbers without mentioning that the *E*-field in Danner et al. was computed on a spherical conductor model and the paper reports the absolute peak value. We use a realistic FEM model with different tissue conductivities and gyral folding, which might produce higher peak cortical fields than a smooth sphere. We take the 99.9th percentile to remove potential artifacts resulting from the FEM. Overall, this mismatch in models might reduce the large margin our stimulator has over the average human motor threshold as reported by Danner et al.

Phosphene thresholds are even less straightforward to estimate. To our knowledge, no study reports an average cortical *E*-field phosphene threshold for monophasic stimulation. The phosphene literature reports thresholds almost entirely in maximum stimulator output (MSO)%. Additionally, the visual cortex sits deeper in the calcarine sulcus, adding significant variability to the analysis. We instead use an MSO% ratio between average motor and phosphene thresholds to estimate whether our stimulator is in the range for effective phosphene induction.

Fidanci et al. [21] found a motor threshold of 50.14 ± 9.75% MSO and a phosphene threshold of 53.93 ± 14.26% MSO. They used a monophasic pulse with a figure-of-eight coil. The effective pulse length was 86 µs, which is very similar to our own device’s effective pulse length of 90 µs. This gives us a phosphene-to-motor output ratio of

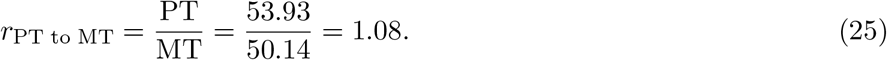

Danner et al.’s conservative estimate places the motor threshold at 63% of our stimulator’s maximum output. Applying the phosphene-to-motor ratio gives us 1.08 × 63% = 68.4% of maximal output, still well within range. The transfer from motor to visual assumes that our coil has a field strength decay over distance similar to that of traditional commercial figure-of-eight coils, which we find to be true when studying the *E*-field decays in SimNIBS of our self-wound coil to the ones from the MagVenture C-B60.

As with motor thresholds, confirmation would require testing on human subjects, which is outside this device’s validated scope.

### 7.7 Validation with clinical TMS

To benchmark our method for deriving the coil’s 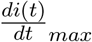 from the pickup-coil-induced voltages, we capture the 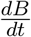 maps of a clinical TMS system. We use the MagVenture MagPro X100 TMS with the MagVenture MRI-B91 coil. We set the MSO% such that the reported `Realized` 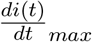 of the stimulator was around the same as our stimulator’s 110.86 A/µs. At 59% MSO, we get a `Realized` 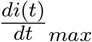 of 110 A/µs.

We proceeded with capturing the 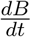 field under the coil using the same methodology as before. During data analysis, a low-pass filter with cutoff frequency *f*_*c*_ = 120 kHz was applied to the data to reduce noise artifacts. We adjusted the modeled coil according to the MagVenture MRI-B91 specifications to ensure the best possible fit.

Our data analysis method estimates the coil 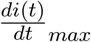 to be 103.89 A/µs with *r*^2^ = 0.9550, which corresponds to a 5.6% error normalized by the stimulator-reported 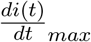. This is similar to the 4.3% error reported by Drakaki et al. when using the same coil and stimulator [19].

Given that the *E*-field is linearly proportional to the 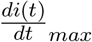, the error we find here transfers to the estimated *E*-fields from §7.6. Even then, our stimulator has significant headroom with respect to the thresholds.

## Ethics statements

This work did not involve human subjects or animal experiments.

## CRediT author statement

**Maxence Lapatrie**: Conceptualization, Methodology, Software, Validation, Formal analysis, Investigation, Writing - Original Draft, Visualization, Project administration, Funding acquisition. **Yuzuha Isetani**: Software, Validation, Investigation, Writing - Original Draft. **Jathav Puvirajan**: Methodology, Validation, Investigation, Writing - Original Draft. **Antonella Catanzaro**: Investigation, Project administration, Funding acquisition. **Siqi Lyu**: Methodology, Validation, Investigation. **Han C. Nguyen**: Methodology, Validation, Formal analysis, Investigation. **William Mathieu**: Validation, Resources, Writing - Review & Editing. **Milica Popovich**: Supervision, Writing - Review & Editing.

## Acknowledgements

This work received financial support from United Builders (https://unitedbuilders.network/).

## Supplementary materials

### High voltage 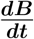 maps

**Figure S1:**
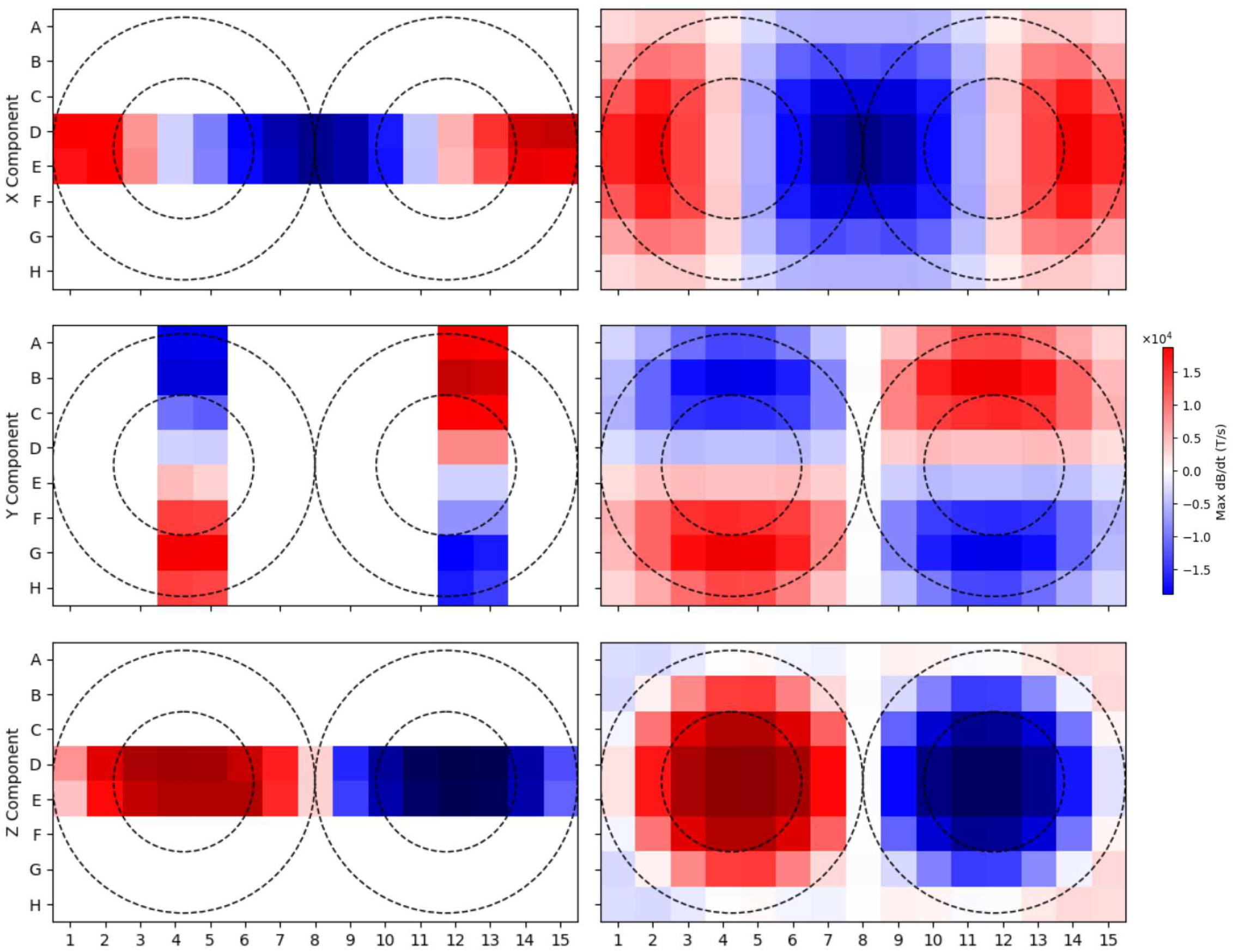
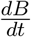 map measured from the presented TMS with the self-wound coil at maximum voltage. The derived map appears on the left, and the theoretical map constructed from the coil model appears on the right.

**Figure S2:**
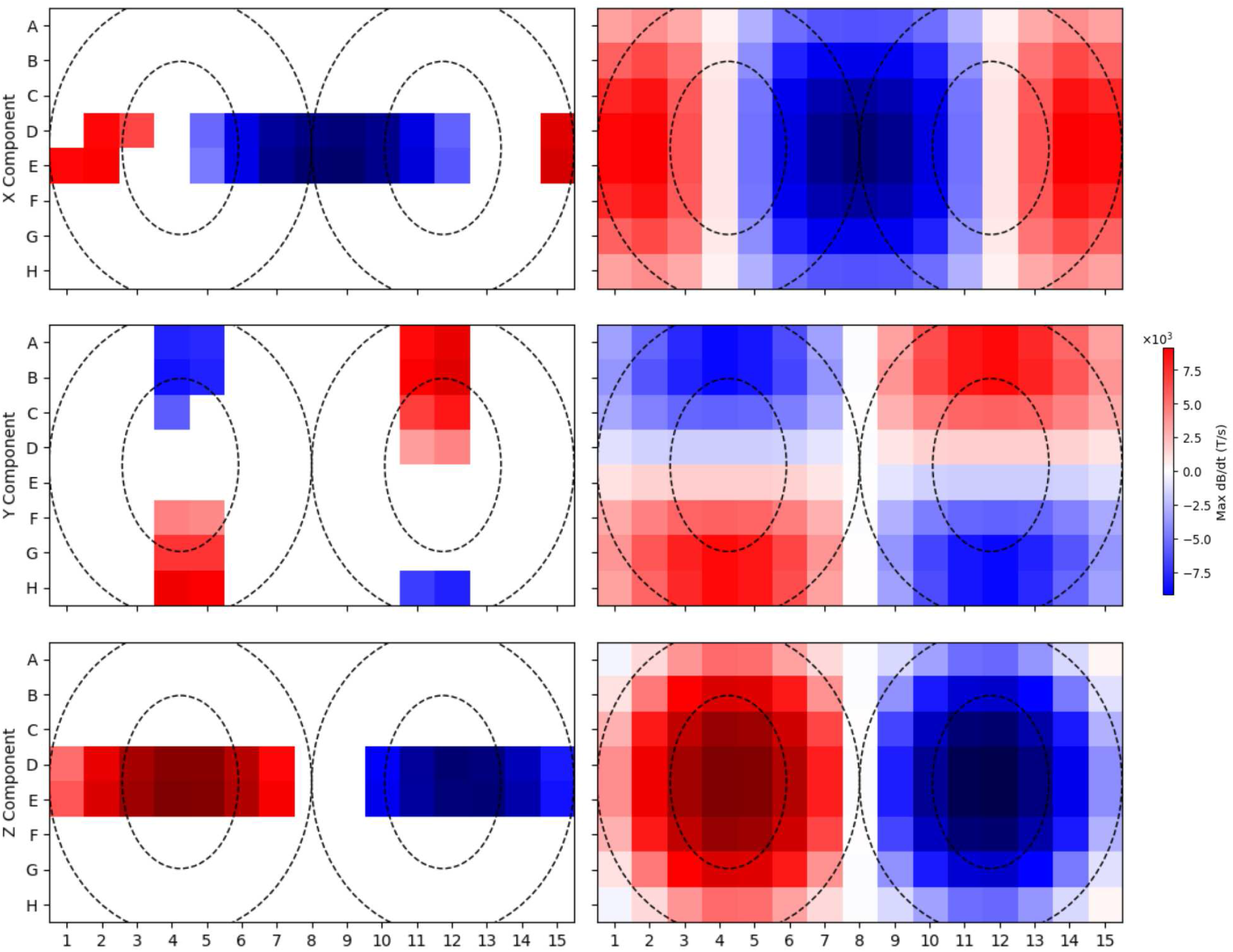
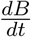 map from the MagVenture MagPro X100 TMS with the MagVenture MRI-B91 coil at a `Realized` 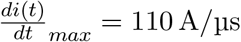. The derived map appears on the left, and the theoretical map constructed from the coil model appears on the right. White squares represent data points rejected for poor signal-to-noise ratio.

### Safety Checklists

#### Device inspection

□ Bank voltage reads 0 V on the connected multimeter.
□ Busbars show no cracking along the flattened edges and have no loosened joints.
□ Nothing is touching the busbars except at the intended connection sites.
□ Bleeders and voltage divider resistors are visually intact and connected at their respective terminals.
□ Safety discharge stick is connected to the bank-negative busbar through discharge resistors and is within reach.
□ Correct transformer tap is selected for the target voltage.
□ Transformer taps are far away from each other.
□ Target voltage does not exceed the system’s maximum voltage.
□ Enclosure lid is closed; lid interlock closed.
□ Dead-man interlock is held by the second person.
□ Comparator trip threshold is set for the intended voltage.
□ Multimeter is readable and enclosed with the high voltage circuitry.
□ PSU current limit set according to expected maximum current draw.
□ TUI divider ratio matches the divider tap actually in use.
□ TUI shows a nominal state. Press R to reset if needed.

#### Pre-charge safety checklist

□ At least two people are present. Both have been briefed on the location and operation of the charge-cutoff switch, the discharge stick, and the de-energization checklist.
□ Operator is wearing Class 1 insulating rubber gloves and insulated footwear. Every person present is wearing impact-rated eye protection.
□ No rings, watches, or other conductive materials on either person.
□ Bench is clear of conductive clutter, tools, and stray wires.
□ No liquids on or near the setup.
□ A physical barrier separates the operator from the high-voltage components.
□ Both people are clear of the high-voltage region and ready for the circuit to be energized.

#### De-energization

□ Stop charging by pressing C again or pressing A to arm the system. The current draw should drop to quiescent levels.
□ Cut the PSU feed either by releasing the dead-man interlock or by pressing the charge-cutoff switch.
□ Turn the PSU off with its respective on/off switch, or by disconnecting the PSU directly from the wall.
□ Verify the multimeter is reading a non-zero voltage. Be wary of zero voltage readings as the multimeter might have been disconnected from the circuit or might be malfunctioning.
□ Allow the bleeders to drain the bank (leave untouched for at least 20 minutes), or
□ Make firm, quick contact with the safety discharge stick hook on the bank positive terminal through the enclosure port. Hold contact for at least 5 seconds.
□ Verify the multimeter reads a bank voltage of 0 V. Verify the multimeter probes are connected to the bank.
□ If the probes of the multimeter appear disconnected, connect the discharge stick for another 10 seconds and leave the device untouched for at least 20 minutes.

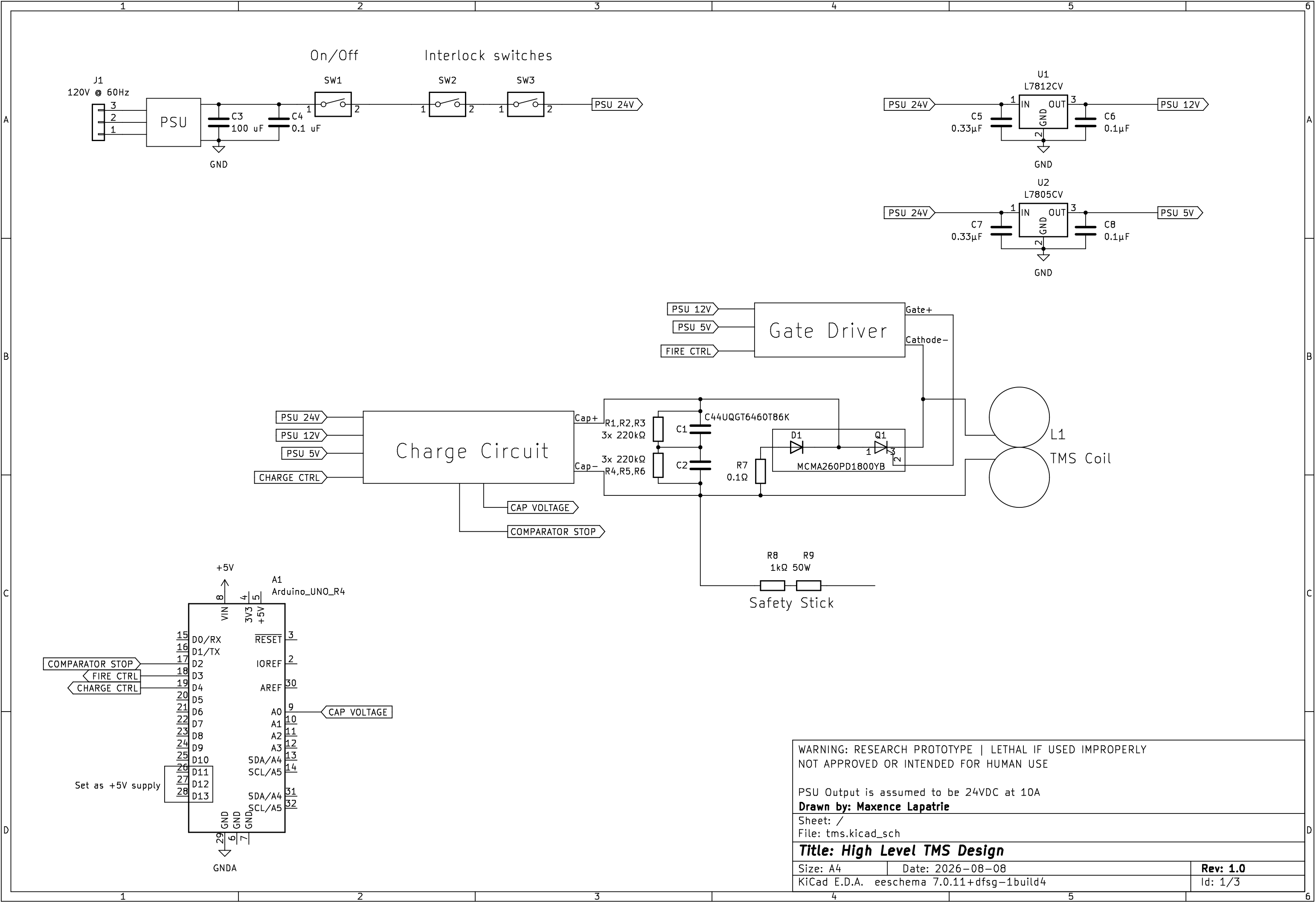

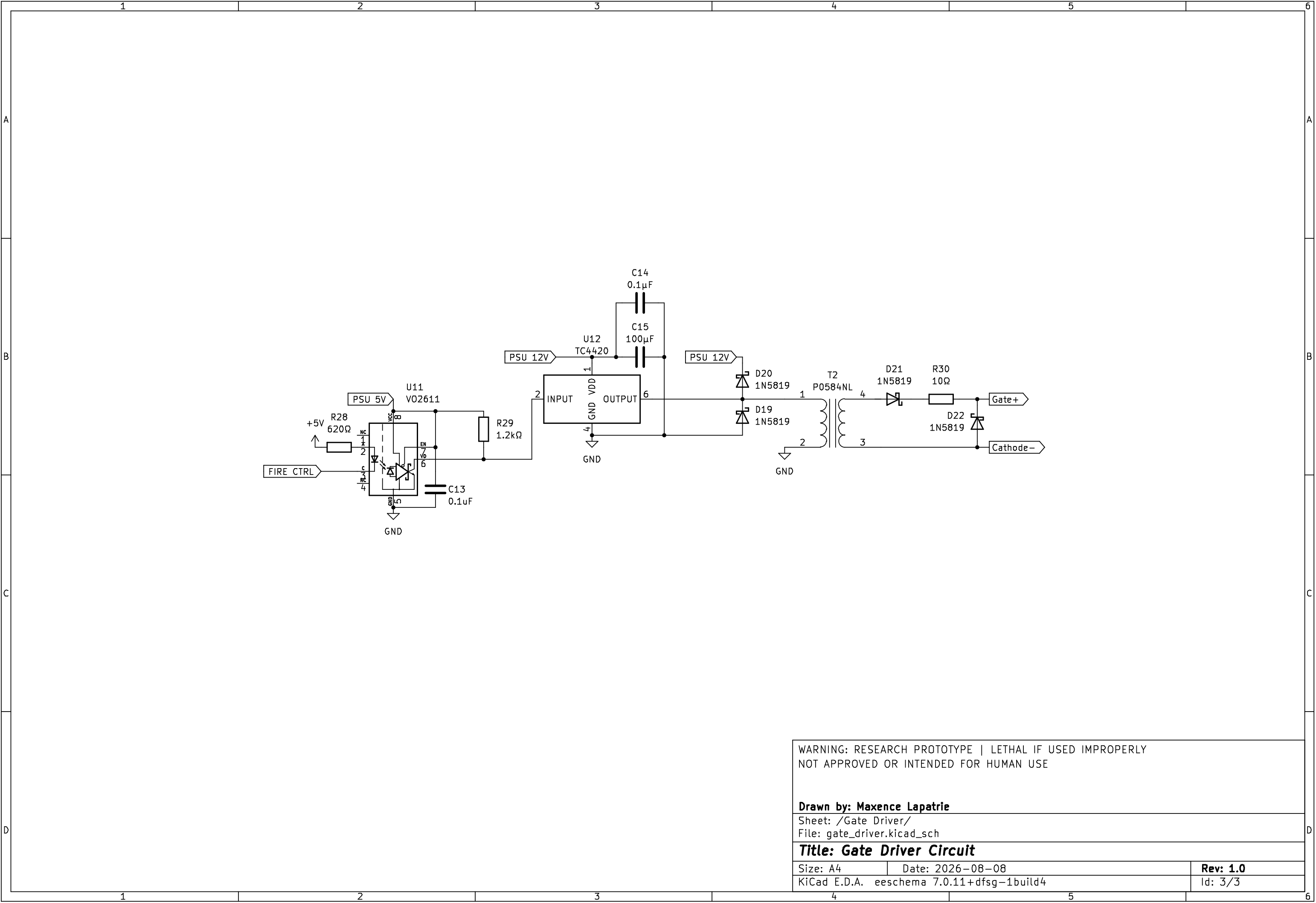

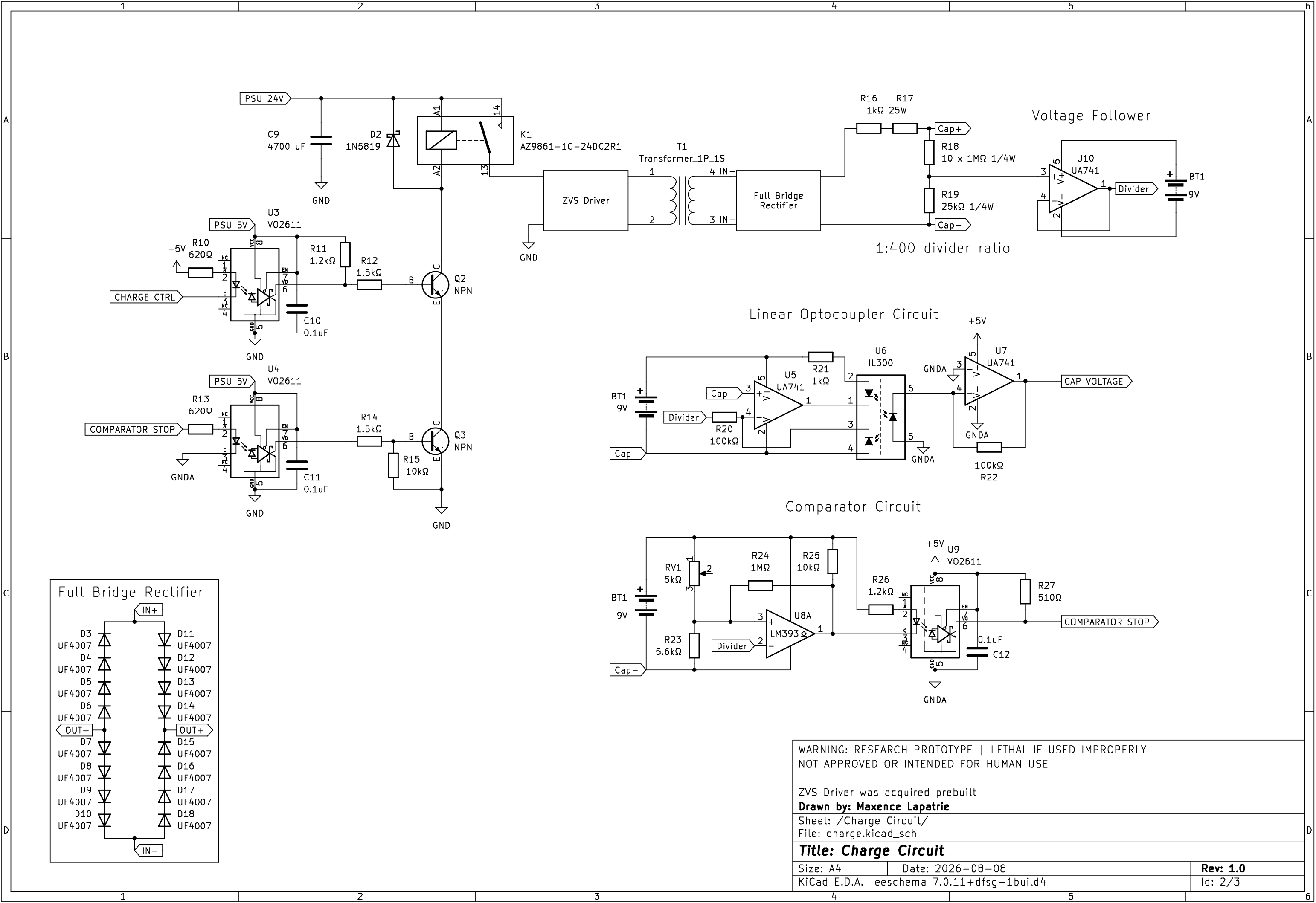

